# Dynamical Diversity in Conductance-Based Neuron Response to kilohertz Electrical Stimulation

**DOI:** 10.1101/2025.08.26.672415

**Authors:** J. G. Polli, F. Kolbl, P. Lanusse, M. G. E. da Luz

## Abstract

Neurons are notably rich in structure and functioning, so rather diverse in their response to stimuli. Consequently, the proper characterization of their dynamical response to external signals is a crucial step in understanding stimulation mechanisms. In particular, kilohertz (kHz) neuronal electrical stimulation tends to induce comportment and drives unseen in (more conventional) lower frequency ranges. Here, we investigate neuronal response of conductance-based models to kHz frequencies stimulation in a broad and often unexplored parameter space region. First, we show that the time evolution exhibited by the paradigmatic Hodgkin-Huxley model under kilohertz stimulation is highly diverse, ranging from regular spiking to chaotic dynamics, as well as displaying regions of complete activity suppression. However, to unveil all these features, a certain level of technical caution is required. For example, we demonstrate that common reductions of sodium dynamics become inaccurate under these frequency regimes. Also, based on suitable markers, we propose a method for mapping the mentioned behaviors on a stimulation parameter space. Second, by extending the study to models of mammalian central nervous system regions, a comprehensive dynamical atlas is obtained. It provides a rather systematic way to typify the response of rapidly forced conductance-based neurons. Thus, the present findings seems to point to an useful scheme for stimulation-based computational neuroscience research at kilohertz frequencies.

## I. INTRODUCTION

Biological neurons constitute paradigmatic examples of complex dynamical systems, comprising large cellular variability interlinked through nonlinear interactions [1]. Indeed, processes occurring at single neuron level give rise to neural circuit phenomena (e.g., rhythmic cycles, synchronization, firing patterns, etc) ultimately emerging in brain functions, such as behavior, memory, and cognition [2, 3]. However, alterations in the structural and functional connectomics of neural circuits may disrupt this emergent complexity, eventually leading to neurological pathologies [4]. In these cases, therapeutic approaches broadly known as neuromodulation [5] rely on chemical, mechanical, and electrical interventions to reorient the dysregulated neural structures. Despite the prevalence of pharmaceutical treatments, its adverse effects [6] and limitations have motivated further developments of the latter type of stimulation.

For example, neuronal electrical stimulation shows clinical efficacy for peripheral and central nervous system’s, PNS and CNS, disorders [7–9]. The usual practice is to deliver stimuli at physiological frequencies, denoted as high-frequencies (HF) [10]. Nonetheless, recent studies with kHz stimulation — called ultra-high-frequencies (UHF) — have led to an effective blocking of abnormal neural propagation [11] and demonstrated enhanced Deep Brain Stimulation (DBS) [12]. Such advances have sparked interest in better characterizing how CNS neurons react to the kHz electrical range [13–17].

Understanding the response of neuronal systems to external forcing has proven a crucial first step in bridging theoretical and clinical therapeutic procedures [18–21]. From a fundamental mathematical perspective, electrical stimulation of neuronal entities can be described as forced nonlinear dynamical systems. Notably, the nonlinear modeling of kilohertz-frequency stimulation in PNS — grounded in the conductance-based Hodgkin-Huxley cable equation approach [22] — have successfully reproduced [23–25] key phenomena observed *in vivo* [26–28]. These models now allow the design of UHF therapeutic protocols for neuromodulation of PNS [29]. Nevertheless, the current literature addressing the response of CNS neurons to UHF kilohertz frequency stimulation remains limited. Experimental evidence revealed the response of hippocampal cells to be diverse [13] and hyperpolarized [14], while Retinal Ganglion Cells (RGCs) presented a correlation between depolarization and spike-suppression both *in vitro* with patch-clamp, and *in silico* (simulation) using point-neuron conductance-based models [15]. Notably, the sole *in-vivo* assessment of UHF neural responses in awake mice pointed to robust cellular and network-level modulation [16]. Spike-suppression traits were evaluated in a point-neuron conductance-based model of Subthalamic Nucleus (STN) [17]. These findings indicate diverse electrophysiological responses to UHF in CNS tissues, potentially associated to different neural scales and thus requiring distinct theoretical descriptions, ranging from point-neurons to network effects. However, a unifying characterization of CNS responses to UHF remains elusive, making a comprehensive investigation of cell-type-specific responses in a broad stimulation parameter space key for elucidating, at least partially, such responses.

In the present study, we investigate the dynamical responses of point-neuron *conductance-based* models for CNS neurons to a breadth UHF stimulation parameter space. In certain contexts [15], this type of modeling has been positively validated in the kilohertz range through direct comparisons with *in vitro* measurements of quantities such as spike-counting and average membrane polarization. This suggests that such a theoretical construction can adequately portray the main aspects of local neuronal response under kilohertz stimulation. Thus, we propose a framework grounded on dynamical systems to characterize and map the UHF-induced dynamics for typical neuromodulation targets; seeking to identify the temporal evolution underlying dynamical neuronal responses to UHF. With this aim, we address structureless neuron models as nonlinear dynamical systems under rapid forcing. The hope is our scheme to help in future computational studies about the response of morphologically more realistic neurons and networks within the UHF regime.

The work is organized as follows. Section II describes the essential of the models (greater technical details are left to the Appendix A). Section III A presents results of trajectories, stroboscopic bifurcation diagrams, Lyapunov exponent, power-spectral densities, and classification of dynamics in stimulation parameter space for the HH model (pertinent methods explanations are given in the Appendix B). Section III B addresses the impact of reducing sodium dynamics to steady state. Section III C develops the dynamical atlas of two retina and five CNS conductance-based neuronal models. Section IV discusses the results and their scope within single-compartmental models as well as comparisons with axonal PNS and perspectives for multi-compartmental setups. Finally, Sec. V drawn the final remarks and conclusion.

## II. A BRIEF OVERVIEW ON THE ANALYZED MODELS

To make the reading easier, here we just outline the general aspects of the addressed models. Much more technical details and explicit expressions are presented in Appendix A.

All the neurons are assumed spatially structureless, Fig. 1(a), with a current clamp represented by the external periodic forcing term

**FIG. 1.**
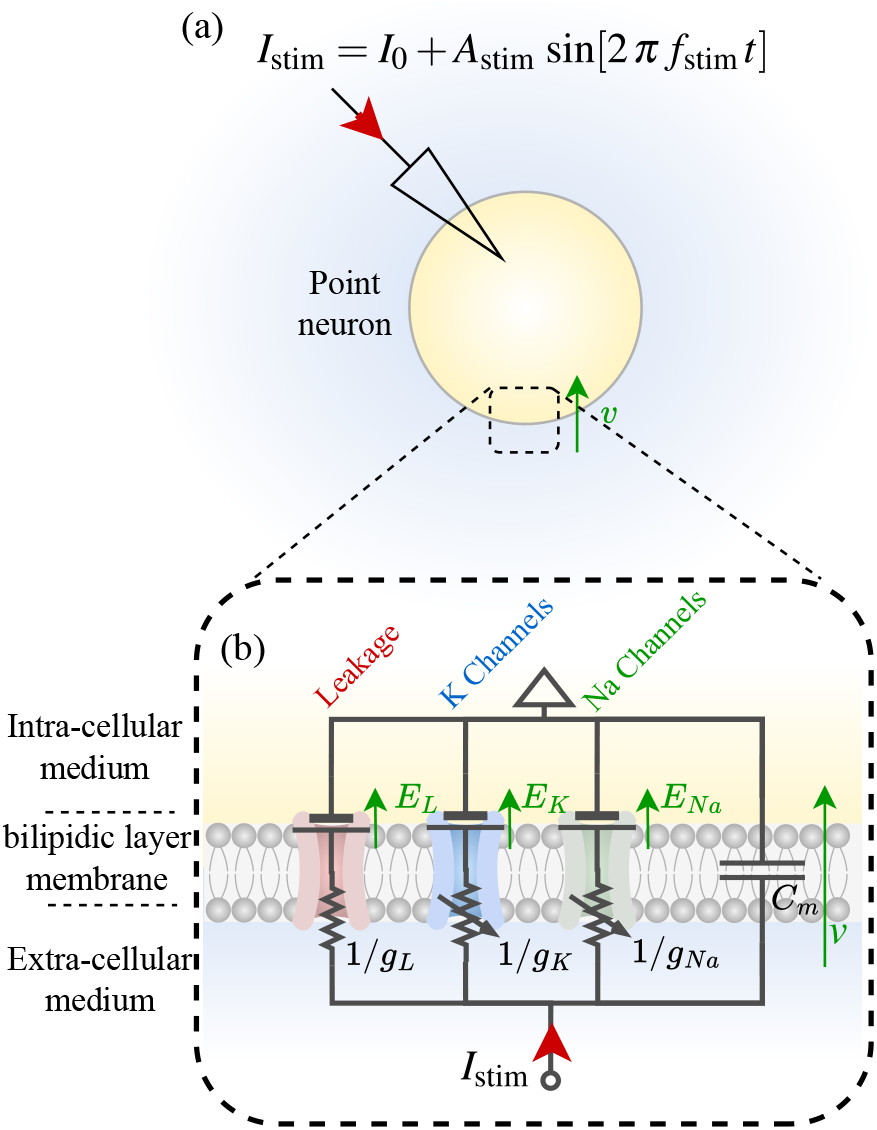
(a) Schematic representation of a point (spatially structureless) neuron stimulated by a current clamp comprising DC and sinusoidal components. (b) Detailed view of the Hodgkin-Huxley (HH) modeling of bilipidic neuronal membrane, where the stimulation current is injected into a conductance-based circuit. The bilipidic neuronal membrane is described as a capacitor of capacitance *C*_*m*_ in parallel with Na^+^, K^+^ and leakage ionic channels, controlled by specific resting potentials and resistors, with dynamical conductances.

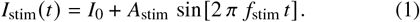

In Eq. (1), *I*_0_ is the direct current (DC) stimulation, whereas *A*_stim_ and *f*_stim_ are the amplitude and frequency of the sinusoidal stimulation. For the UHF response, we focus on supraphysiological regime, namely, 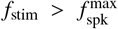, for 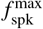 the maximum firing-rate achieved under DC stimulation, i.e., *I*_stim_ = *I*_0_ (refer to Fig. 2(a) and the discussions in Sec. III A).

**FIG. 2.**
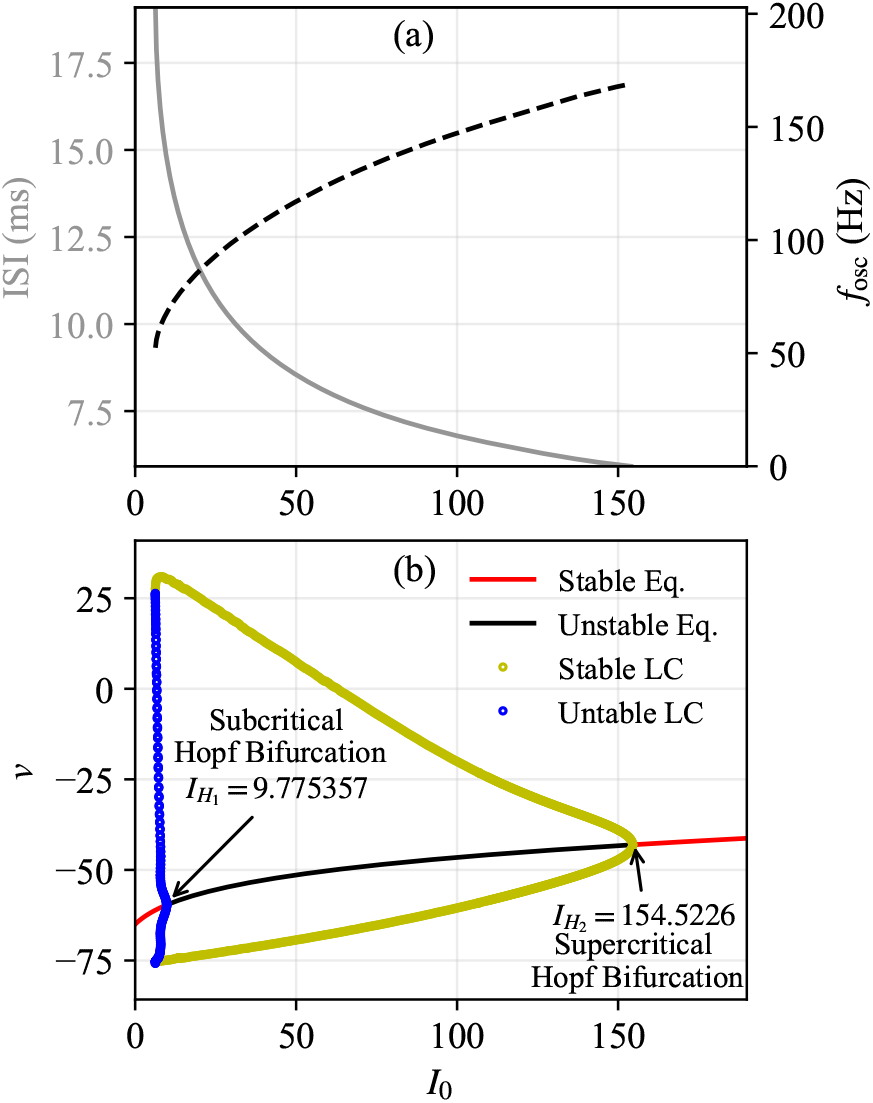
(a) Inter-spike interval (ISI) and spiking frequency *f*_osc_ = 1/ISI, illustrating the class-II behavior of the HH model (i.e., displaying a positive minimal frequency). (b) Bifurcation diagram for the *I*_0_ parameter, exhibiting stable and unstable equilibrium as well as stable and unstable limit cycles (LC). Their stability is shifted by Hopf bifurcations (subcritical 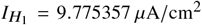 and supercritical 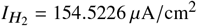).

We first consider the Hodgkin-Huxley (HH) model [22] for probing supraphysiological neuron response. The HH, a conductance-based dynamical system of the giant squid axon, describes the membrane potential *v* (*t*) via a nonlinear system of ordinary differential equations. Its equivalent circuit, Fig. 1(b), represents the neuronal membrane as a capacitor — with membrane capacitance per unit of surface area *C*_*m*_ (in *μ*F/cm^2^) — which is in parallel with three resistive branches. These are ionic channels, related to leakage (L), potassium (K), and sodium (Na) currents, with respective voltage-dependent conductances *g*_*L*_, *g*_*K*_, *g*_*N a*_ and constant reversal potentials *E*_*L*_, *E*_*K*_, *E*_*N a*_. Further details are given in the Appendix A 1.

The HH model has been extensively explored, displaying relevant dynamical trends like bifurcation phenomena under DC clamp [30–33]. Also, under periodic forcing at physiological frequencies, the HH model has shown subharmonic locking, quasi-periodicity, chaos [34–36], and advanced coding properties for more complex inputs [37]. However, at UHF studies are considerably limited [15, 17], not comprehensively exploring the proper parameter space. Therefore, we discuss such a model for a large portion of amplitude-kilohertz-frequency stimulation parameter, investigating the resulting trajectories, the stroboscopic bifurcation map, spectral components, Lyapunov exponents, and systematically identifying distinct dynamical regimes.

A fundamental technical aspect, likewise important for the understanding of neural functioning is the following. The HH model is four-dimensional, of variables *v, m* (sodium activation) and *h* (inactivation) and *n* (potassium) gating. At physiological frequency regimes, the system can be reduced to three or even two dimensions [38–40] through simplification or elimination of the sodium dynamical variables. However, full investigations about the effects of such a reduction in faster stimulation regimes, to the best of our knowledge, are lacking. In special, sodium channels play a critical role in the propagation block mechanism observed in PNS under UHF regimes [41, 42]. In this way, for our complete analysis of the HH, we solve the full model, and then run careful comparisons with the reduced versions.

We second address generalizations, proposed in the literature, of the standard HH, discussing the response of conductance-based models of CNS mammalian under kilohertz frequency stimulation. These include cortical [43], hippocampal (pyramidal excitatory [44] and basket inhibitory cells [45]), subthalamic nucleus (STN), globus pallidus (GPe) [46], and retinal ganglion cells (RGCs) [47]. The main assumptions and dynamical equations of all these HH extensions are summarized in Appendix A 2, in particular refer to Table I. The modeling of the aforementioned regions have shown a diverse range of traits for periodic forcing at physiological frequencies [48–51]. Nevertheless, for kilohertz stimulation, no systematic survey of the response parameter space has been performed. By investigating Lyapunov, trajectory, and parameter-space comportment in the same manner as for the HH model, we examine the map of emergent dynamics in the supraphysiological stimulation parameter space. Our findings quantify cell-type attractors and provide tools for global parameter exploration, including hitherto unexplored regions.

**TABLE I.**
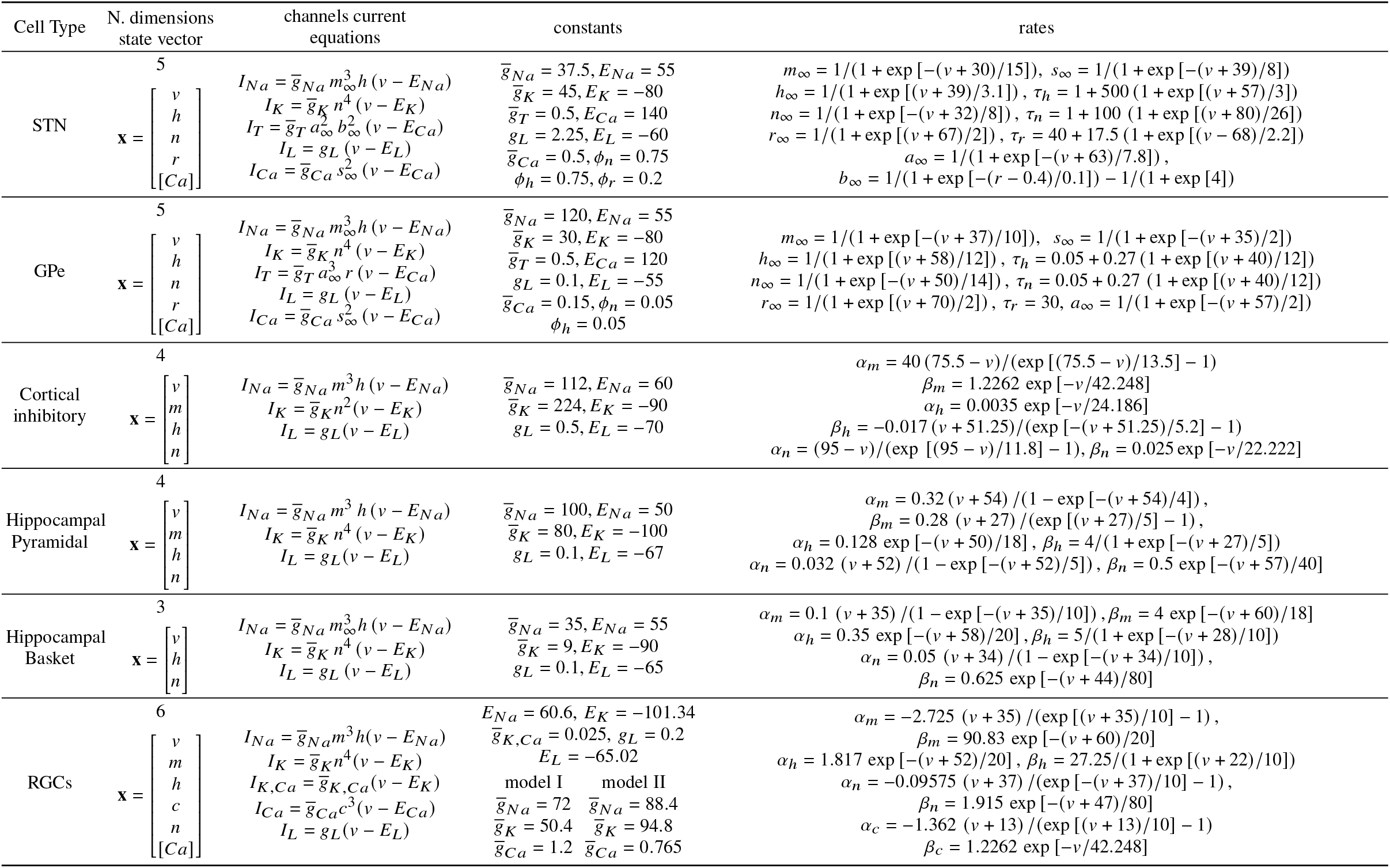
The key equations and parameter values for the mammalian neuronal models considered in this work. The units are suppressed, but the conductance is in mS/cm^2^, reverse potentials in mV and time constants in ms.

## III. RESULTS

### A. The HH model diverse dynamical behaviors in the stimulation parameter space

In the following we address the response of the HH model to kilohertz electrical stimulation, examining the resulting trajectories and establishing a framework for frequency–amplitude parameter space exploration. But prior to that, we briefly review some known trends of the HH model under constant stimulation, i.e., *A*_stim_ = 0, with *I*_stim_ = *I*_0_ for a varying *I*_0_.

In Fig. 2(a) we show the evolution of Inter Spike Interval (ISI) and frequency of oscillation (*f*_osc_ = 1/*I SI*) as *I*_0_ increases. In Fig. 2(b), we also show the bifurcation diagram with the stability of equilibrium and limit cycles for the extrema of the membrane potential *v* at each value of *I*_0_. We note that between subcritical and critical Hopf bifurcations, a stable limit cycle with positive minimal frequency is observed. This characterizes the HH neuron model as of class-II [52], so in opposition to class-I, where neurons fire at arbitrarily small frequencies. Also, for the HH neuron model, the maximum attained frequency is 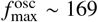 Hz.

#### 1. Trajectories and spiking regimes

For sake of comparison with prior works [17], for a given stimulation frequency *f*_stim_ and amplitude *A*_stim_ we can define the quantity

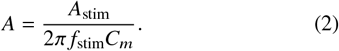

Physically, *A* (expressed in mV) represents the amplitude of induced voltage in a purely capacitive membrane.

Consider now *A*_stim_ > 0 and 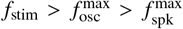. Figure 3(a)-(d) display examples of *v* (*t*) for a neuron forced with *I*_0_ = 10.0 *μ*A/cm^2^ and stimulation frequency *f*_stim_ = 1.0 kHz, for different amplitudes *A*_stim_. For reference, under constant input *I*_0_ = 10 *μ*A/cm^2^ (*A*_stim_ = 0 *μ*A/cm^2^) the equilibrium is unstable, leading to periodic spiking with a period *T* = 14.6 ms and spiking frequency of *f*_osc_ = 68.3 Hz.

**FIG. 3.**
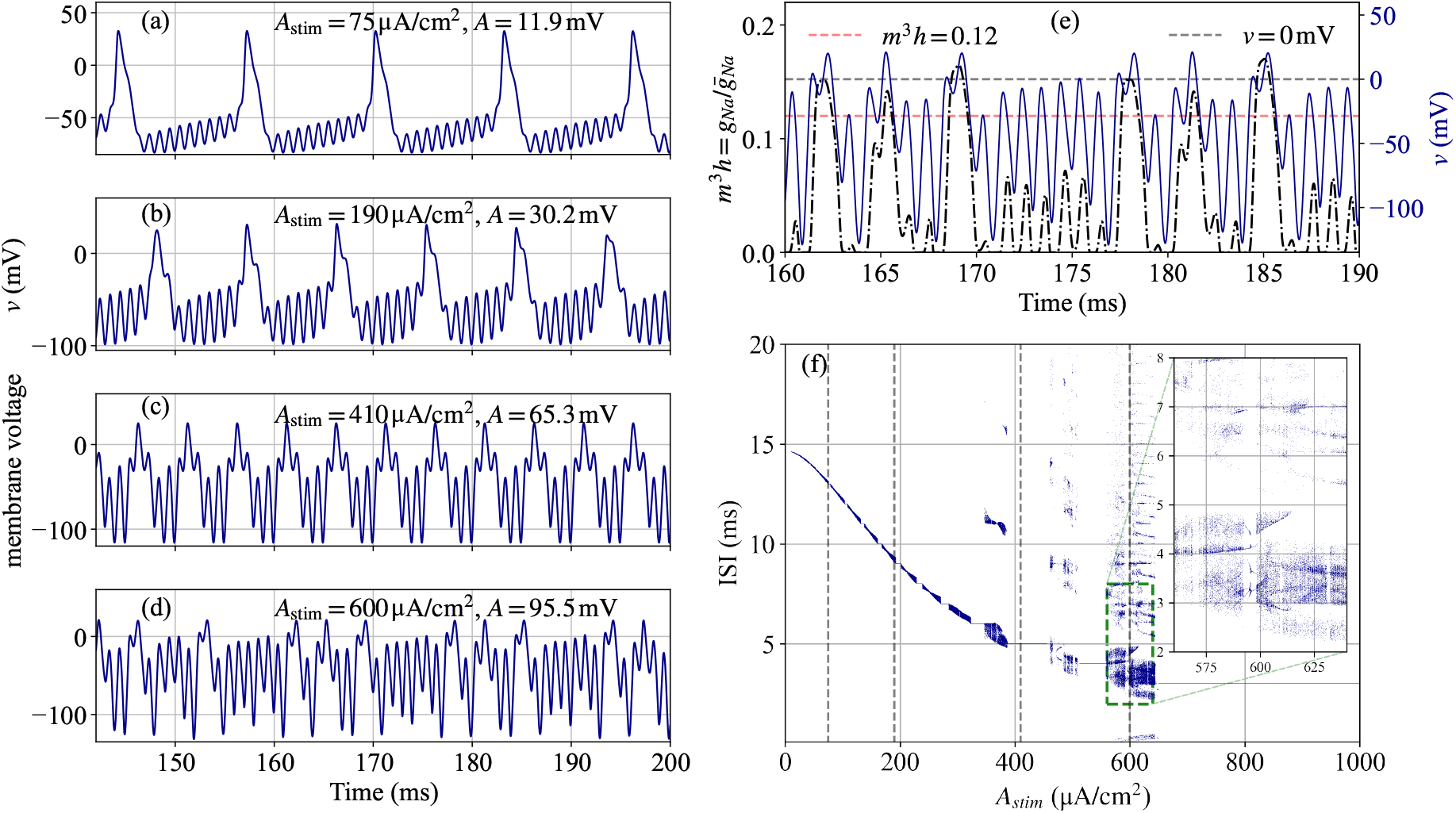
Membrane potential time series, *v* (*t*), and ISI (inter-spike interval) for the HH model stimulated with *I*_0_ = 10.0 *μ*A/cm^2^, *f*_stim_ = 1.0 kHz and representative values of *A*_stim_. (a) The case *A*_stim_ = 75 *μ*A/cm^2^ shows a periodic spiking behavior. (b) When *A*_stim_ = 190 *μ*A/cm^2^, the spiking dynamics ceases to be periodic. (c) *A*_stim_ = 410 *μ*A/cm^2^ leads to a periodic trajectory with ISI 5 ~ ms, whereas (d) *A*_stim_ = 600 *μ*A/cm^2^ displays an aperiodic spiking dynamics. (e) The sodium relative conductance 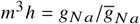 along with the trajectory for *v* (*t*) for *A*_stim_ as in (d). The spiking thresholds *m*^3^*h* = 0.12 and widely used *v* = 0 are indicated as references. (f) The ISI for *m*^3^*h* = 0.12 taking as spiking threshold and the values of *A*_stim_ ranging from 10 *μ*A/cm^2^ to 1000 *μ*A/cm^2^. The gray dashed lines marks the *A*_stim_ value for the trajectories in (a)-(d). The inset zooms up a region where the ISI is far from periodic.

Figure 3(a) shows trajectories for stimulation with amplitude *A*_stim_ = 75 *μ*A/cm^2^. From Eq. (2), this yields a voltage amplitude *A* ≈ 11.9 mV. For this strength, the high-frequency components only modulate the underlying low-frequency spiking, consistently with averaging model predictions [17]. Figure 3(b) shows the dynamics for a higher stimulation amplitude, *A*_stim_ = 190 *μ*A/cm^2^ (*A* ≈ 30.2 mV). This elevates the spiking frequency — a trend potentially extrapolated from the DC stimulation bifurcation diagram (see Fig. 2). However, individual spikes now exhibit clear distortion from the forcing waveform. The resulting evolution no longer resembles DC stimulation behavior, nor locks harmonically with the stimulation frequency, although maintaining repetitive patterns. Notably, this dynamics matches *in vitro* experimental observations of RGCs under kilohertz-frequency stimulation [15]. Further, Fig. 3(c)-(d) depicts dynamics at even higher stimulation amplitudes: *A*_stim_ = 410 *μ*A/cm^2^ (*A* ≈ 65.3 mV) and *A*_stim_ = 600 *μ*A/cm^2^ (*A* ≈ 95.5 mV), respectively. In Fig. 3(c), periodic spiking reemerges but with increased frequency and stronger modulation compared to Fig. 3(a). In Fig. 3(d), prominent spikes disappear, replaced by irregular oscillations with variable refractory periods. This variability aligns with known kilohertz-frequency stimulation effects [10]. Moreover, the trajectory irregularity resembles chaotic dynamics, a trend with desynchronization properties in networks [53]. We later quantify these qualitative observations employing Lyapunov analysis.

Periodicity in neural dynamics can be typified by inter-spike interval (ISI) measurements, requiring precise spike timing detection. However, as shown in Fig. 3(a)-(d) external forcing artifacts often difficult spike identification in membrane potential *v* traces. Thus, we use the relative sodium conductance 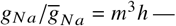 responsible for initializing depolarization in the HH model — as a reliable spike indicator (see Appendix A 1 for conductance details). In Fig. 3(e) we show an example of how the relative sodium conductance (left axis) compares with the membrane potential (right axis) for same stimulation parameters as in Fig. 3(d). While the membrane potential undergoes an unpredictable oscillatory comportment, the sodium relative conductance clearly distinguish active/inactive cellular states. So, we set a common conductance-based spiking threshold *m*^3^*h* = 0.12 (red dashed line in Fig. 3(e)), registering a spike when the relative conductance surpasses this value from bellow. Actually, this is more reliable than to set membrane potential spiking thresholds (e.g. *v* = 0 mV, gray dashed line) once such a usual procedure frequently registers double-crossings during forced oscillations.

Figure 3(f) displays a representative ISI diagram for *f*_stim_ = 1 kHz and stimulation ranging from 10 to 1000 *μ*A/cm^2^. Vetical gray dashed lines mark the *A*_stim_ values of Fig. 3(a)-(d), ordered from left to right. For *A*_stim_ = 75 *μ*A/cm^2^ and *A*_stim_ = 410 *μ*A/cm^2^, a constant ISI indicates purely periodic behavior. For *A*_stim_ = 190 *μ*A/cm^2^ a narrow band of ISI values suggests quasi-periodicity. Within 10–350 *μ*A/cm^2^, quasiperiodic and periodic regions alternate. Beyond 350 *μ*A/cm^2^, ISI values distribute across multiple bands, presenting periodic windows that take turns with more complex patterns. For instance, at *A*_stim_ = 600 *μ*A/cm^2^, ISIs range widely from 1 ms to 20 ms. A blow up into a region around *A*_stim_ = 560 and *A*_stim_ = 640 *μ*A/cm^2^ reveals complex ISI distributions, including bifurcation cascades and shading — characteristic of chaotic trajectories.

#### 2. Stroboscopic orbits diagram: classification of observable dynamics

To inspect a wide range of possible dynamics for the forced HH neuron model, one can construct stroboscopic orbit diagrams by sampling trajectories at every *T*_strob_ = 1/*f*_stim_ time intervals. Figure 4(a) shows such orbits for parameters *f*_stim_ = 1.25 kHz, *I*_0_ = 0*μ*A/cm^2^ and stimulation amplitude ranging from 0 to 3000 *μ*A/cm^2^. In the plots, background color highlight different ranges of spiking dynamics. Light lavender indicates absence of spikes (*A*_stim_ < 140.5 *μ*A/cm^2^), while light red identifies spiking regions (*A*_stim_ > 140.5 *μ*A/cm^2^). Within a spiking region we have periodic stroboscopic dynamics (i.e. subharmonic of the forcing frequency), where *P*_*k*_ denotes a *k*-cycle stroboscopic orbit. As *A*_stim_ increases, the system shifts from *P*_1_ (no-spike region) to a regime that alternates between aperiodic and periodic windows. Notably, within the spiking zone, periodic sectors appear in a sequence of decreasing periods {…, *P*_5_, *P*_4_, *P*_3_, *P*_2_}. Extra examples of bifurcation diagrams are presented in the Supplemental Material (SM), Fig. S1 and Sec. S1.

**FIG. 4.**
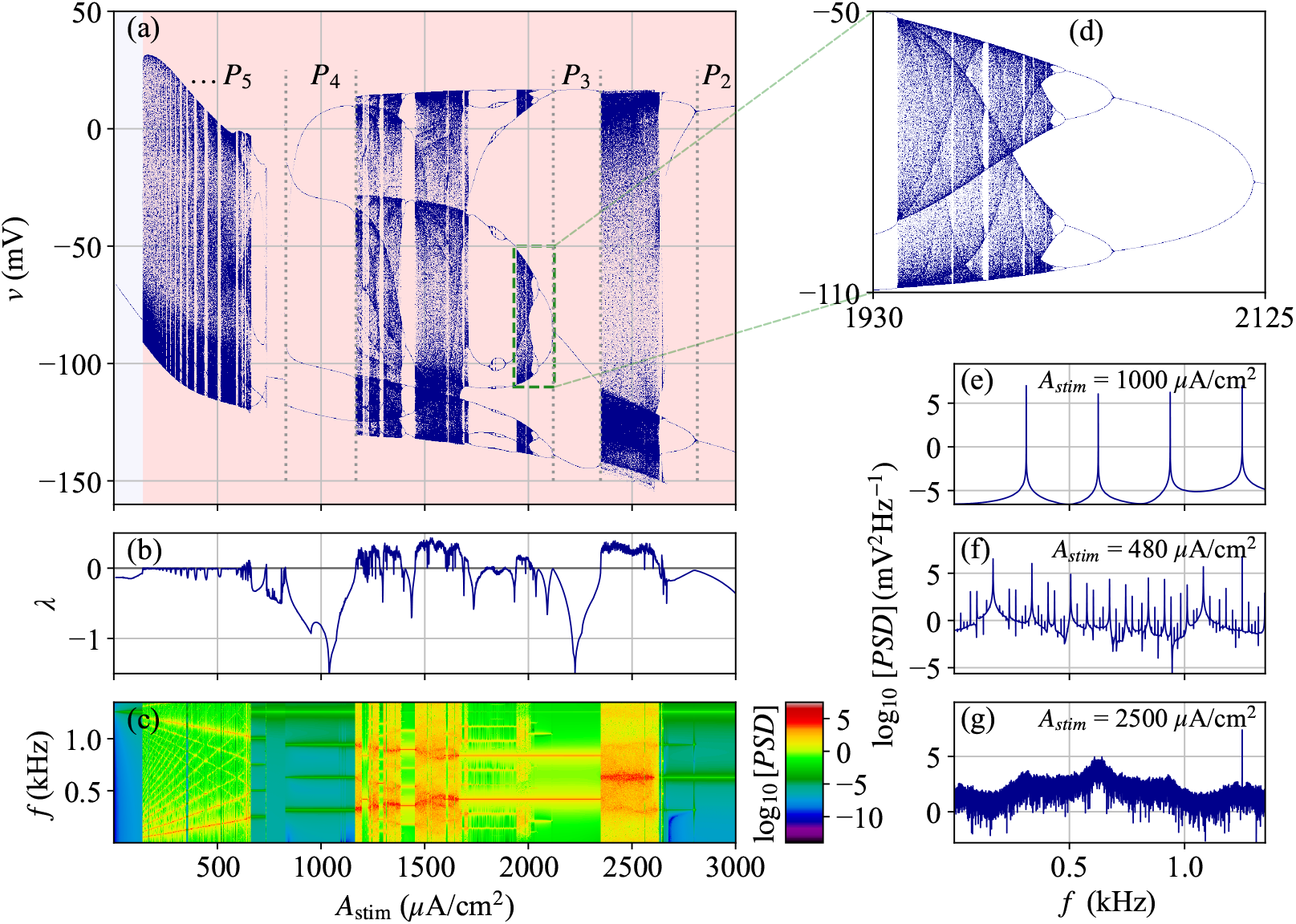
Dynamical transitions and spectral features of the kilohertz-frequency stimulated HH neural model for *f*_stim_ = 1.25 kHz and *I*_0_ = 0 *μ*A/cm^2^. (a) Stroboscopic orbits diagram of the membrane potential *v* versus stimulation amplitude *A*_stim_ ∈ [10, 3000] *μ*A/cm^2^. Background color indicates no-spiking (white) and spiking (light red) dynamics. Vertical dotted lines delimit examples of subharmonic zones (*P*_*k*_ for *k* the stroboscopic period). (b) Maximum Lyapunov exponent *λ* for the same parameter range. (c) Logarithm PSD showing distribution of spectral components. (d) A blow up of the stroboscopic orbit diagram of (a) in the interval *A*_stim_ ∈ [1930, 2125] *μ*A/cm^2^ showing a reverse period-doubling bifurcation route to chaos. Spectral profiles for three regimes, (e) quasi-periodic (*λ* ~ 0) for *A*_stim_ = 500 *μ*A/cm^2^, (f) *P*_4_ subharmonic for *A*_stim_ = 1000 *μ*A/cm^2^, (g) chaotic (*λ* > 0) for *A*_stim_ = 2500 *μ*A/cm^2^.

In Fig. 4(b) we depict the corresponding maximum Lyapunov exponent (MLE) *λ* (see Appendix B) for the trajectories in Fig. 4(a). The values are strictly negative before reaching the no-spike/spiking transition, indicating stable periodic trajectories. Along the transition, *λ* switches between nearzero and strictly negative values. A more detailed inspection reveals near-zero *λ* matching the aperiodic regions in the stroboscopic orbits diagram, enabling quantitative classification of these quasi-periodic (QP) trajectories. Positive *λ*’s, pointing to chaotic dynamics, start around *A*_stim_ = 625 *μ*A/cm^2^. Chaotic zones (*λ* > 0) alternate with periodic windows until the last observed aperiodic sector, before the *P*_2_ region. To clearly distinguish periodic and aperiodic regimes, we computed the power spectral density (PSD) of trajectories (see Appendix B for details). Fig. 4(c) presents the PSD results for the trajectories in Fig. 4(a). A prominent line at *f* = 1.25 kHz corresponds to the forcing frequency. Notice that for all amplitudes, the forcing frequency acts as a modulation in the associated dynamics. Periodic regions *P*_*k*_ exhibit *k* distinct subharmonic peaks.

Figure 4(d) shows a zoom into the red-dashed rectangle in Fig. 4(a), detailing a reverse period-doubling cascade, well known to be a route to chaos [54]. This is consistent with the positive *λ* values of this region in Fig. 4(b). Figure 4(e) represents the spectral profile for *P*_4_ periodic dynamics, where 4 distinct peaks are observed in the PSD. Quasi-periodic zones (*λ* ~ 0) display multiple sharp dominant frequencies superimposed on the background frequencies, as exemplified in Fig. 4(f). In contrast, chaotic regimes (*λ* > 0) show broadband noise without dominant peaks. This is illustrated in Fig. 4(g), where besides the forcing frequency, no distinct peaks emerge above the background noise.

Figure 4 is computed for *I*_0_ = 0 *μ*A/cm^2^ and *f*_stim_ = 1.25 kHz. In Sec. S1 of the SM, we display stroboscopic bifurcation diagrams for additional combinations of DC currents, *I*_0_ = 0, 5, 10 *μ*A/cm^2^, and stimulation in the kilohertz range, i.e., *f*_stim_ = 1, 1.25, 2.75, 5 kHz. We observe that the amplitude value at which there is a transition (from no-spike to spiking) decreases as *I*_0_ increases. If 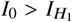, the no-spike phase disappears, as expected from the analysis in Fig. 2.

In total, we found four distinct responses of the HH model to kilohertz frequency in the frequency–amplitude parameter space. (i) Non-spiking (NS) dynamics: taking place below a threshold stimulation amplitude, 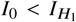. (ii) Periodic sub-harmonic dynamics: membrane potential oscillations phase-lock with a subharmonic of *f*_stim_, exhibiting constant spiking frequency, period-*k* (*P*_*k*_) stroboscopic dynamics, *λ* < 0, and *k* subharmonic peaks in the PSD. (iii) Quasi-periodic (QP) dynamics: aperiodic trajectories with *λ* ~ 0 that bridge consecutive *P*_*k*_ and *P*_*k*−1_ periodic windows. (iv) Chaotic dynamics: displaying aperiodic trajectories with *λ* > 0. It emerges primarily through period-doubling cascades.

In fact, a fifth comportment also occurs with non-zero DC current, *I*_0_ ≠ 0 *μ*A/cm^2^. For constant stimulation, the HH model exhibits periodic oscillatory behavior between Hopf bifurcations 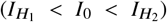. Figure 5 shows a stroboscopic orbits diagram for 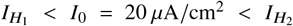 and *f*_stim_ = 5 kHz. A first region with multiple spikes can be observed until a threshold of *A*_stim_ = 497 *μ*A/cm^2^. For *A*_stim_ > 497 *μ*A/cm^2^, the membrane voltage turns into a non-spiking *P*_1_ behavior, with values typical of hyperpolarization. As discussed in [17], we call this trend spike-suppression (SS), since the application of UHF represses neural activity (we mention we have reproduced the result in [17] with identical parameters). Importantly, SS differs from NS. The DC current alone would elicit spiking, but here the sinusoidal stimulation acts towards suppressing the spikes.

**FIG. 5.**
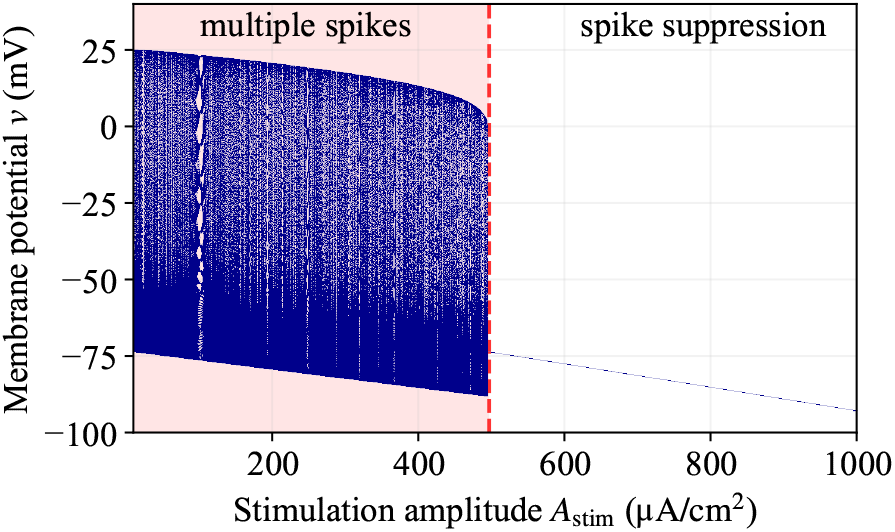
Stroboscopic plot for *f*_stim_ = 5 kHz and *I*_0_ = 20 *μ*A/cm^2^, illustrating spike-suppression as *A*_stim_ increases. At small amplitudes, spikes are present until a threshold (here of 497 *μ*A/cm^2^). Beyond it, no spike is triggered and the membrane potential tends to decrease fairly linearly with *A*_stim_, indicating that the cell is hyperpolarized.

#### 3. Dynamical mapping of the parameter-space

For the kilohertz-frequency range, we next qualitatively characterize the frequency-amplitude parameter space. In Fig. 6(a) we plot the MLE *λ* for *A*_stim_ ∈ [0, 7000] *μ*A/cm^2^, *f*_stim_ ∈ [0.4, 3.5] kHz and DC current *I*_0_ = 0. The exponent *λ* was calculated only for *A* < 475 mV (see Eq. (2)), thus avoiding unphysiological hyperpolarization regimes. Chaotic regions (*λ* > 0) occupy dense bands for stimulation frequencies below 3 kHz and amplitudes from few tens to about 6500 *μ*A/cm^2^. Moreover, these bands arise within 40 mV < *A* < 425 mV, demarcated by red (40 mV) and yellow (425 mV) dashed lines in Fig. 6(a).

**FIG. 6.**
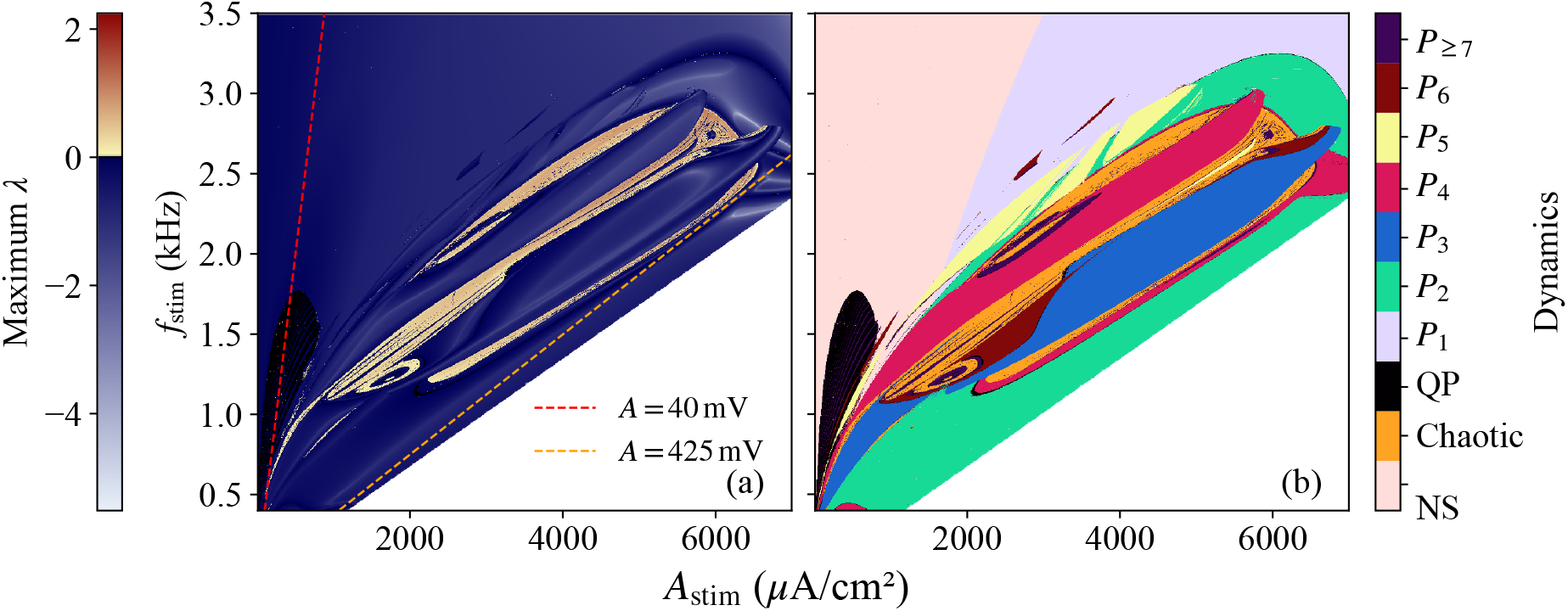
(a) MLE map of the HH model for *f*_stim_ ∈ [0.4, 3.5] kHz, *A*_stim_ ∈ [100, 7000] *μ*A/cm^2^ and DC current *I*_0_ = 0. Trajectories with *A* > 475 mV yield non-physiological hyperpolarization (in addition being hard to numerically integrate). They correspond to the lower left part of the plots (white triangles). Red and yellow dashed lines delimit the region between 40 mV < *A* < 425 mV. (b) Functional mapping of the HH model for the same parameter space region of (a). It displays regions of non-spiking (NS), chaotic, quasi-periodic (QP) and subharmonic of periods *P*_*k*=1,…,6_ as well as *P*_*k*≥7_. The algorithm to obtain the different regions is explained in the Appendix B 4.

We further classify the response of the HH model to each frequency-amplitude combination using an algorithm described in Appendix B 4. It identifies the five behaviors specified in Sec. III A 2, namely, chaotic, quasi-periodic (QP), periodic (*P*_*k*_), no-spiking (NS) and spike-suppression (SS). Figure 6(b) shows the resulting dynamical map for *I*_0_ = 0 (related to Fig. 6(a)). As previously described, chaotic zones form bands. QP zones are limited to a region in the bottom-left part of the map, vanishing above 1.75 kHz and amplitude ~ 1000 *μ*A/cm^2^. NS are present in the top-left map region, corresponding to lower induced membrane voltage amplitudes. SS is absent once *I*_0_ = 0 prevents spiking without sinusoidal forcing. Periodic regions are scattered over a great fraction of the parameter space, with higher-period orbits embedded within QP and chaotic zones. All periodic regions tend to *P*_2_ as the induced membrane potential amplitude *A* increases, leading to a *P*_2_ dominance for *A* > 475 mV. Moreover, transitions are observed between distinct regimes (in the functional mapping, Fig. 6(b)): QP ↔ NS (bottom-left), NS ↔ *P*_*k*_ (top left), QP ↔ *P*_*k*_ (bottom left), *P*_*k*_ ↔ *P* _*j*_ (*j* ≠ *k*) and *P*_*k*_ ↔ chaotic (center).

At this point, a brief summary of the previous results is in order. UHF stimulation induces diverse dynamical regimes in the HH model. Although the distribution of these regimes in the parameter space is somehow involved, extended regions of uniform behavior are present. The density and dimensionality of the chaotic and periodic regions are still open questions (and a problem which hopelly we will address in a forthcoming contribution). Nonetheless, they are relevant for targeting specific neuronal responses in neural engineering procedures, see the Discussion section.

### B. Supra-physiologic stimulation effects on sodium channels dynamics

The mechanism of UHF stimulation in the PNS primarily involves sodium channel blocking, although the possibility of targeting potassium channel has not been fully excluded in the literature [42]. For CNS stimulation, the impact of supraphysiological frequencies on ionic channels remains elusive. Conductance-based models could, at least as a first approximation, clarify this key issue by determining how UHF stimulation modulates the dynamics of gating variables in ionic transport. Thus, here we explore the action of UHF on dimension reductions of the HH model, where certain gating variables are removed.

As detailed in the Appendix A 1, the original 4D HH model — specifying *v* (*t*), *m* (*t*), *h* (*t*) and *n* (*t*) — admits two distinct three-dimensional reductions. (1) The first by replacing the *m*_∞_ variable by its steady-state *m* value. (2) And the second by eliminating the *h* variable by supposing a linear relation between *h* and *n*. A third, 2D reduction, which we call (3) combines (1) and (2). Since the Na conductance depends on the quantity *m*^3^ *h*, the aforementioned simplifications directly impact on sodium dynamics. However, these simplifications are widely used since they have a mildly influence on the model dynamics at low frequencies. In (1)–(3), the bifurcation analysis for the DC stimulation (*A*_stim_ = 0 *μ*A/cm^2^) preserves the succession of subcritical and supercritical Hopf bifurcation points. Indeed, this leads to the same dynamical comportment presented by the full 4D HH model, only changing the 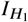 and 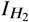 values (see SM Section S2 for details).

Figure 7(a-c) show stroboscopic bifurcation diagrams for the full 4D, the 3D *h* reduced and the 3D *m* reduced HH models, respectively. All share the same forcing frequency (*f*_stim_ = 1 kHz) and stimulation amplitude range (*A*_stim_ ∈ [0, 700] *μ*A/cm^2^). The systems exhibit a progression of periodic and quasi-periodic trends as *A*_stim_ increases. Remarkably, the 3D *h* reduced closely matches the sequence of periodic/quasiperiodic regimes of the 4D model, with significant deviations emerging only at high amplitudes. In contrast, the 3D *m* reduced model displays important distinctions for these sequences, being much more sparse and displaying a different stimulation amplitude range.

**FIG. 7.**
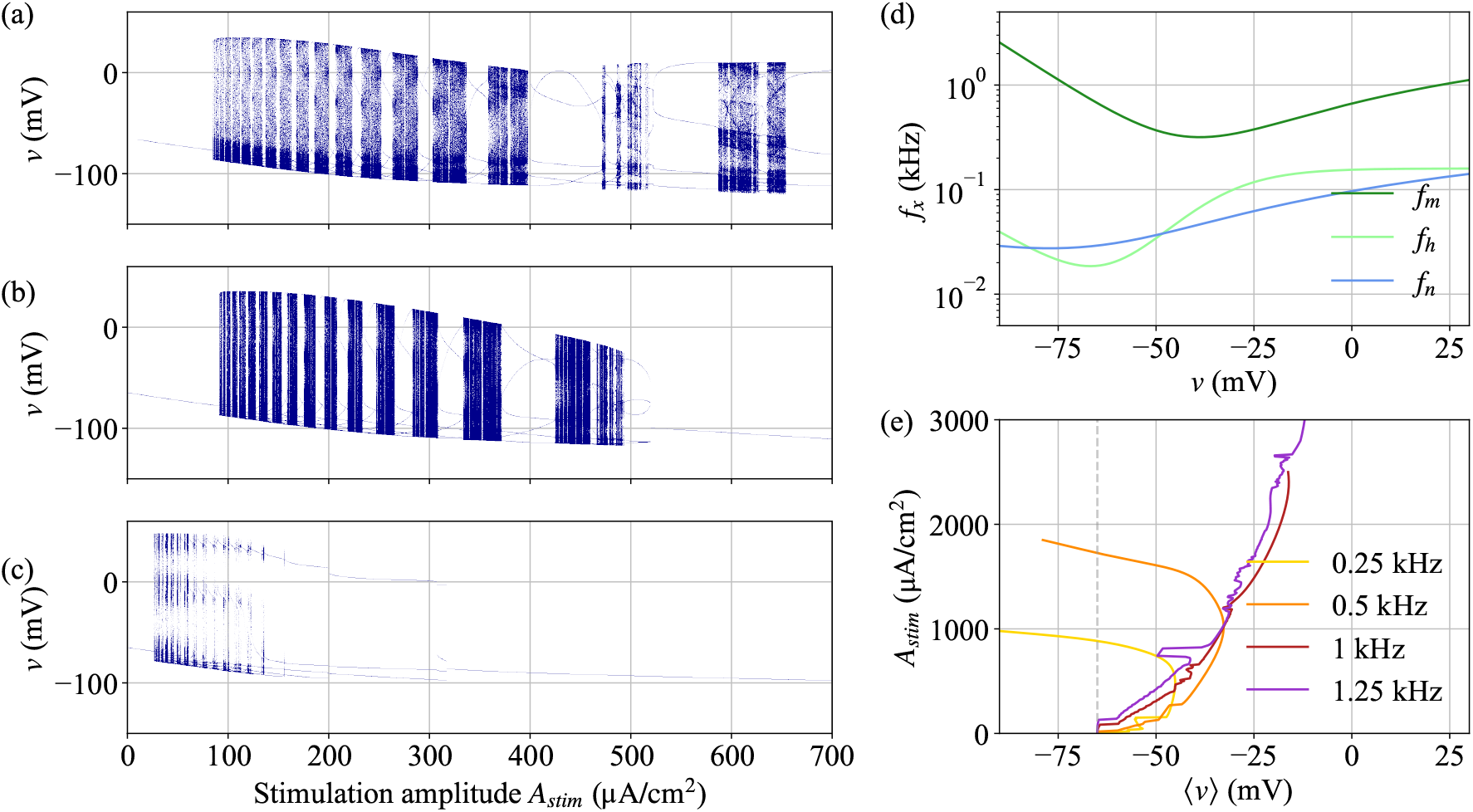
Stroboscopic, frequency domain and baseline analyzes for 4D and 3D HH models. Stroboscopic bifurcation diagrams are shown for the full 4D, (b) 3D *h* reduced and (c) 3D *m* reduced cases. In all them the forced frequency is *f*_stim_ = 1 kHz, the DC *I*_0_ = 0 and the stimulation amplitude *A*_stim_ ∈ [0, 700] *μ*A/cm^2^. Compared to the original 4D model, the 3D *h* maintains a similar structure of periodic/quasiperiodic successions, while for the 3D *m* most of the quasi-periodic and chaotic-like orbits disappear. (d) The gating variable transition frequency *f*_*x*_ (with *x* being *m, h* and *n*) as function of *v* for the 4D HH model. Both *h* and *n* attain to the physiological range of frequencies, but with *m* displaying transition frequencies in the supra-physiological 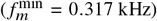. (e) The stimulation amplitudes versus the averaged baseline membrane potential ⟨*v*⟩ for four values of the stimulation frequencies for the 4D HH model. The gray dashed line denotes the resting potential *v* = − 65 mV, such that baselines to its left (right) are hyperpolarized (depolarized). For *f*_stim_ = 0.25 and 0.5 kHz the baseline shows initial depolarization but becomes hyperpolarized as *A*_stim_ increases. For *f*_stim_ = 1.0 and 1.25 kHz, only depolarization is observed.

Chaotic regimes (*λ* > 0) take place in both 3D models (see SM, Figs. S5 and S6). In the *h* reduced case, chaos arises for *A*_stim_ in the interval 480–500 *μ*A/cm^2^. For the *m* reduced version, the chaotic region emerges in a narrow window around *A*_stim_ = 135 *μ*A/cm^2^. The progression of periodic regimes shows fundamental differences among models. The 4D displays the characteristic sequence … *P*_5_, *P*_4_, *P*_3_, *P*_2_ for higher stimulation amplitudes. In contrast, the sequence is truncated in the 3D models. For the *h* reduced, *P*_5_ transitions directly to SS, while for the *m* reduced, the transition is from *P*_4_ to SS.

The 2D model dynamics (see SM, Fig. S7) somewhat resembles that of 3D *m* reduced model, but with two relevant differences. First, chaotic regions are absent, which is consistent with the expectations for systems of dimension *d* < 3. Second, the transitions occur from the periodic regime *P*_3_ to SS.

For each gating variable *x*, either, *m, h* or *n*, we have (see Eq. (A6) in Appendix A 1)

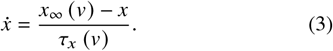

As given in Eq. (A5), *x*_∞_ (*v*) and *τ*_*x*_ (*v*) represent the voltage-dependent *x* steady state value and the *x* associated time constant. These quantities are related to sigmoidal-shaped rate functions *α*_*x*_ (*v*) and *β*_*x*_ (*v*). Our findings demonstrate that the membrane voltage *v* presents a non-trivial spectral response to UHF (see Fig. 4(d)). Moreover, because the steady-states *x*_∞_ (*v*) are monotonic functions of *v*, they inherit spectral components of *v*, including dominant components around *f*_stim_.

The Eq. (3) further allows to interpret *x*_∞_ (*v*) as a forcing term for the *x*-gating variable dynamics. Additionally, it describes a low-pass filter, characterized by a voltage-dependent transition frequency (in Hertz)

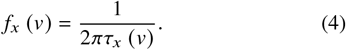

Assuming that *f*_stim_ dominates the spectrum of *x*_∞_, then the gating variable exhibit two distinct regimes. When *f*_stim_ ≪ *f*_*x*_, the dynamics of *x* (*t*) closely follows *x*_∞_ (*v*), so that *x* ≈ *x*_∞_ (*v*) is a rather good approximation. Conversely, when *f*_stim_ ≳ *f*_*x*_, the dynamics of *x* (*t*) cannot track the external forcing, hence *x* ≈ *x*_∞_ (*v*) is inaccurate. Such a regime has been explored for suitable stimulation amplitudes using classical average theory [17, 55] — see Sec. IV C for a detailed discussion.

For the 4D HH model, Fig. 7(d) depicts the evolution of the transition frequencies, Eq. (4), for typical values of *v*. For the *h* and *n* gating variables, the transition frequencies lie in the range of tens to hundreds of Hz. Therefore, under both HF (~ 130 Hz) or UHF stimulation, these variables operate in the regime *f*_stim_ ≳ *f*_*x*_, invalidating assumptions like *h* = *h*_∞_ and *n* = *n*_∞_. In contrast, the transition frequency 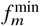 associated with the *m* gating variable spans from around 317 Hz up to few kHz. Thus, for HF stimulation (or even lower) the simplification *m* = *m*_∞_ works well. But for kilohertz stimulation, the condition *f*_*m*_ ≪ *f*_stim_ no longer holds, invalidating the 4D to 3D reduction approach. Indeed, as shown by the stroboscopic bifurcation plots in Fig. 7(c), the 3D *m* model fails to reproduce the characteristic dynamical structure of the full 4D HH under kilohertz stimulation. In this way, one can conclude that the influence of supra-physiological forcing is directly linked to sodium channels, and in particular to their activation dynamics, governed by the *m* gating variable.

Figure 7 (e) shows the average membrane potential ⟨*v*⟩ (see Appendix B) across a range of stimulation amplitudes and four different stimulation frequencies (*f*_stim_ = 0.25, 0.5, 1.0, 1.25 kHz) for the original 4D model. Such quantity, also referred to as *baseline*, has been modeled and experimentally measured in neural cells [15]. Our results align with previous findings: (i) for lower stimulation frequencies (*f*_stim_ = 0.25 kHz and 0.5 kHz), the baseline initially increases (depolarizes) at low amplitudes, then decreases toward resting potential and eventually hyperpolarizes as the amplitude increases; and (ii) for larger stimulation frequencies (*f*_stim_ = 1.0 kHz and 1.25 kHz), the baseline monotonically depolarizes with amplitude. Remarkably, (ii) corresponds to stimulation frequencies way above the minimal transition frequency of the *m* gating variable, reinforcing the role of the Na activation dynamics in shaping the response to supra-physiological stimulation.

Lastly, Fig. 8 presents a stroboscopic bifurcation diagram in which *f*_stim_ increases while the induced membrane potential amplitude *A* is held constant so that *A*_stim_ varies accordingly, refer to Eq. (2). When 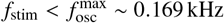 (blue dashed line in Fig. 8), the system exhibits a *P*_1_ periodic response. This behavior remains unchanged as *f*_stim_ crosses both 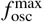 and 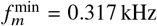 (green dashed line in Fig. 8), with the neuron continuing to phase-lock with the stimulation. As the frequency is further increased, a sequence of transitions takes place: from *P*_1_ to *P*_4_, getting back to *P*_2_, followed by QP dynamics, then *P*_3_ and next going through a cascade of period doubling bifurcations, culminating into chaotic behavior. Although this does not account for the whole range of observed behaviors, it suggests that phenomena associated with UHF arise not only when 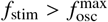, but also for 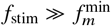.

**FIG. 8.**
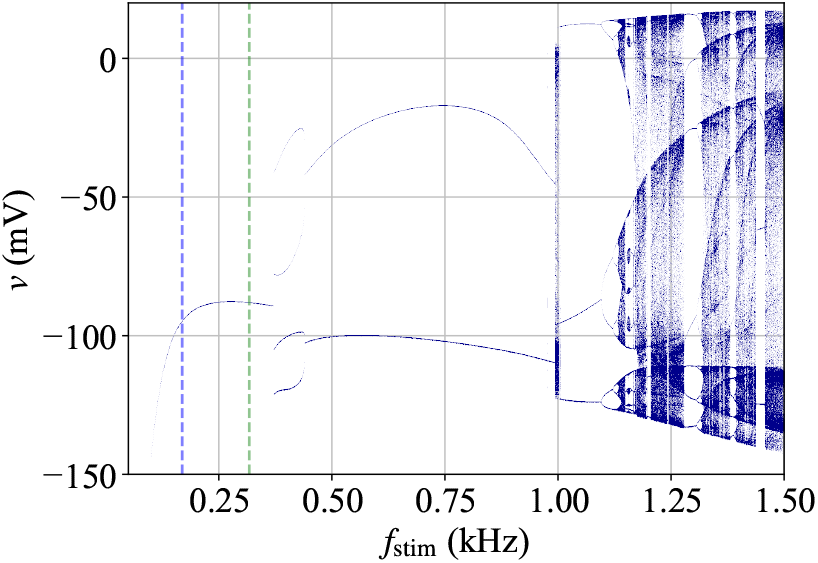
Stroboscopic plot of *v* for *f*_stim_ ∈ [0.15, 1.5] kHz and a varying *A*_stim_ accordingly to maintain constant the induced voltage amplitude *A* ≈ 191 mV, Eq. (2). It shows the neural behavior when crossing the physiologic/supra-physiologic limit (blue dashed line at 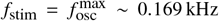) and the frequency for which the *m* gating variable transition frequency is minimal (green dashed line at 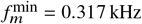).

### C. Supra-physiologic stimulation effects on distinct mammalian neural models

The HH model has been extensively investigated for over fifty years, serving as a valuable neuron prototype to support and even benchmark more realistic studies (see, e.g., [17]). However, it was derived based on non-mammalian (squid) axon data, so not capturing the dynamical diversity of different brain structures. Therefore, to explore the effects of UHF in mammalian brain — trying to be in line with certain known important processes [15, 17] — we performed a mapping of behaviors induced by kilohertz stimulation discussing a collection of models aimed to distinct brain regions.

Concretely, next we address models for: (1) cortex [43], (2) hippocampus pyramidal excitatory cells [44], (3) hippocampus basket inhibitory cells [45], (4) subthalamic nucleus (STN) [46], (5) globus pallidus external (GPe) [46], and retinal ganglion cells (RGC) [47] split into two, (6) small cellular area (model I) and (7) large cellular area (model II). All them are formulated under the nonlinear conductance-based scope of the HH model, but in certain cases including new specific currents (see Appendix A 2). For DC stimulation, such models can be classified either as Type I (continuous firing-rate) or Type II (discontinuous firing-rate) [52], as detailed in SM Section S5.

Figure 9 shows dynamical mapping for mammalian neural models, following the same procedure of Fig. 6(b) (refer to Appendix B 4). All mappings are generated assuming a constant DC current *I*_0_ = 10 *μ*A/cm^2^. The stimulation frequency *f*_stim_, vertical axis, ranges are in the regime 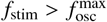, for 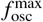 the (model-specific) maximum oscillatory frequency, which also exceeds the corresponding model maximum spiking frequency 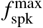. These thresholds are computed via bifurcation analysis under DC conditions and given in SM Tab. S2 (type I models) and Tab. S3 (type II models). Maximum frequency and amplitude intervals are chosen so to capture the full diversity of dynamical trends for each model.

**FIG. 9.**
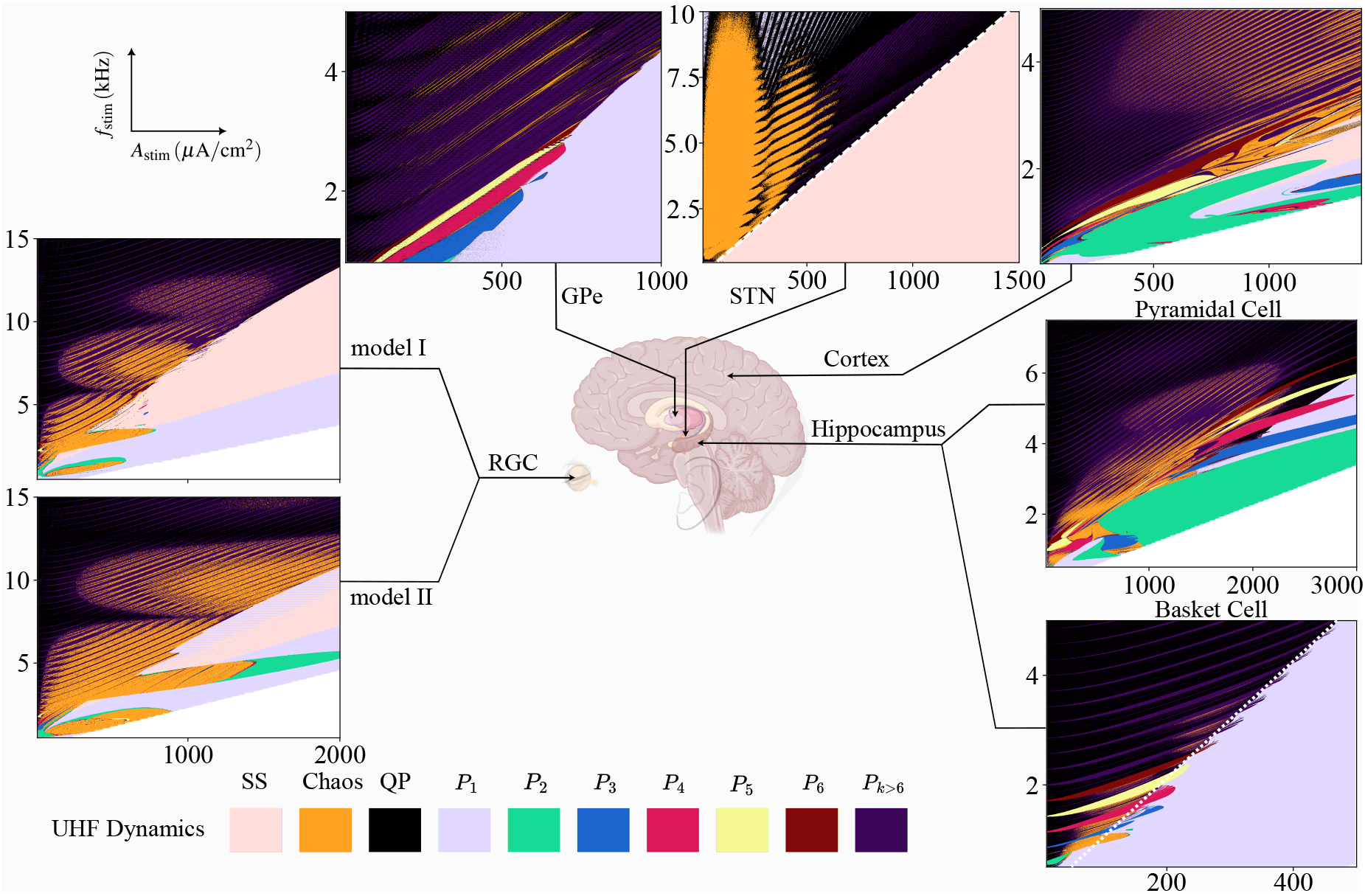
Mapping of the UHF-induced stimuli dynamics of various conductance-based models in the amplitude-frequency parameter space. They address the structure of mammalian brain of: Retina (RGC), external Globus-Pallidus (GPe), SubThalamic Nucleus (STN), Cortex and Hippocampus Piramidal, and Basket. For all cases, the DC current is *I*_0_ = 10 *μ*A/cm^2^. The horizontal (vertical) axis displays the range of stimulation amplitude *A*_stim_ in *μ*A/cm^2^ (stimulation frequency *f*_stim_ in kHz). Each color represents a different dynamical behavior, namely, non-spiking (NS), chaos, quasiperiodic (QP), phase-locked (*P*_1_) and sub-harmonics *P*_*k*_ for *k* = 2, 3, 4, 5, 6 as well as higher order sub-harmonics *P*_*k*>6_. White dashed (dotted) line in STN (hippocampal basket cell) map represents constant *A* = 23 mV (*A* = 15.1 mV).

For such a DC current of *I*_0_ = 10 *μ*A/cm^2^, all models present spike at *A*_stim_ = 0. In this way, regimes for which *A*_stim_ > 0 without the presence of spiking are classified as spike-suppression (SS). Actually, SS regimes are observed in both RGC, STN, and cortical cell, but are absent in GPe and in the two hippocampal cell models. For RGCs and STN, SS occupies well-defined large portions of the parameter space, while for the cortex, it appears in a small area at the central right part of the associated map in Fig. 9. Chaotic zones arise in all cases, although they are confined to small regions in GPe and hippocampal basket models. QP and sub-harmonics (*P*_*k*_) are ubiquitous, especially in the upper-left regions of the parameters space (high frequency, low stimulation), where induced membrane potential *A* values are small.

Significantly, the functional mapping across models is highly diverse, with no universal pattern describing the location or arrangement of dynamical regimes. Moreover, none of the maps in Fig. 9 display any pattern similar to the original HH model in Fig. 6 (b). Below we detail the specific features of each model parameter space.

Models for Cortex and Hippocampus (pyramidal and basket cells) exhibit qualitative similar responses in their parameter spaces. In all cases, the top-left region (high frequency, low amplitude) is dominated by alternating QP and high period subharmonic behavior (*P*_*k*>6_). In cortical and pyramidal cell models, the central portion of the parameter space lacks a clear structure, alternating between chaotic, sub-harmonics of low period (*P*_1_–*P*_5_) and SS. As the induced membrane potential *A* increases (which is equivalent to moving toward the lower right corner of the plots), both models lock into the dynamics *P*_2_, similar to the usual HH. In contrast, basket hippocampal cells present a more organized structure in the parameters space central region, with alternating bands of chaotic and periodic regions. Further, as the induced membrane potential *A* is increased, a large *P*_1_ region emerges (light blue, below the white-doted line which marks *A* ~ 15.1 mV).

Despite being governed by the same conductance model (with different parameter sets, see Tab. I), GPe and STN display contrasting dynamical responses. The STN presents large chaotic, QP, and periodic regions, whereas the GPe predominantly shows periodic regions (computed up to *P*_200_), with only small QP and chaotic bands. In the STN map, a clear boundary separates spiking from SS regions, indicated by a white dashed line (along which *A* ~ 23 mV). In the GPe, distinct transitions between periodic regimes are observed, nevertheless the frontiers are not characterized by clear constant values for *A*.

The RGC models I and II lead to similar structures, typified by intercalated QP and high-period subharmonic regimes at high-frequency and low stimulation amplitudes (upper-left part of the parameter space), resembling the dynamics of the cortical and hippocampal models. At lower frequencies (1 kHz < *f*_stim_ < 10 kHz) chaotic regions emerge embedded within periodic regimes. A large SS region occupies the central-to-upper-right portion of the parameter space, transitioning into the *P*_1_ regime as the induced membrane potential *A* increases (lower-right). Although some details of our stimulation amplitude differ from [15], the occurrence of chaotic, periodic and QP comportment at low *A*_stim_’s is consistent with the spike-rich regime reported in [15]. Additionally, the mid region of spike suppression concurs with the suppression described in such work. The correlation between SS regions and depolarization effects is further considered in Section IV A.

Importantly, unless for GPe and Basket Hippocampus, all the other models in Fig. 9 exhibit (in the parameter space) the full collection of dynamical regimes identified in the previous sections, namely, SS, chaotic, quasi-periodic, and subharmonic *P*_*k*_. Although non-spiking is absent in Fig. 9, it does arise when *I*_0_ is below the spike onset threshold (see SM, Fig. S15), hence similar to Fig. 6 for the HH model. Some models present either SS or *P*_1_ for high induced membrane potential *A* values, both behavior representing pure harmonics of the external forcing. However, they should be assumed distinct regimes, once the spiking threshold condition *m*^3^*h* > 0.12 is not achieved for SS. Also, if this threshold is changed, certain SS and *P*_1_ regions may interchange.

We should mention that the basket hippocampal, STN and GPe models models assume the simplification *m* = *m*_∞_ (*v*). As emphasized in Section III B, this reduction might be inaccurate when *f*_stim_ ≳ *f*_*m*_. Transition frequencies *f*_*x*_, Eq. (4), for the gating variables of mammalian models are given in SM, Fig. S14. For these basket hippocampal model, *f*_*m*_ ranges from several hertz to a few kilohertz. Then, under kilohertz stimulation *f*_stim_ ≳ *f*_*m*_ the assumption *m* = *m*_∞_ (*v*) is inaccurate, eventually resulting in loss of dynamical variability and shifts in the true locations of dynamical regimes in parameter space. For STN and GPe, *f*_*m*_ (*v*) is not defined (refer to [46]), preventing extra analysis and comparison with *f*_stim_. But these models play a significant role in computational studies of DBS for Parkinson Disease (PD). Thus, our findings suggest that sodium activation dynamics should be added to STN and GPe models to investigate kilohertz DBS.

## IV. DISCUSSION

The previous general analyzes aimed to properly and comprehensively characterize the phenomenological behavior of the full HH model as well as of its mammalian extensions under UHF stimulation. These are relevant theoretical constructions to describe distinct types of point (so reduced) neurons. In the remainder of this contribution, we shall discuss the bio-physiological consequences of our results, pinpoint how our study could be useful for future computational investigations of morphologically accurate neurons — e.g., with the inclusion of spatial diffusion terms to qualitatively represent axons — and networks going through kilohertz stimulation.

### A. Correlation between depolarization and spike-suppression

In different analyses, membrane depolarization has been correlated with spike suppression in RGC by means of single and multi-compartmental neuron simulations [15, 56, 57]. This suggests that modeling based on point reductions of the neuronal morphology can, at least for patch-clamp experiments, properly describe actual neurons local response to UHF. Nevertheless, our results show that SS represents only one of many dynamical responses observed in the models. Therefore, next we shall discuss how other dynamical regimes relate with membrane potential polarizations.

Figure 10 presents the average membrane potential variation, ⟨Δ*v*⟩ = ⟨*v*⟩ − *v*_0_, where ⟨*v*⟩ is the time-averaged membrane potential after filtering the external signal artifact and *v*_0_ is the baseline before stimulation onset, see Appendix B. Figure 10 (a) and (b) depict ⟨Δ*v*⟩ for the RGC I and STN, respectively. The DC current is *I*_0_ = 10 *μ*A/cm^2^ (see SM Fig. S16 for HH and mammalian models). In the RGC model I, depolarized regions (⟨Δ*v*⟩ > 0) qualitatively match the SS regions in the functional map of Fig. 9. Nonetheless, weaker levels of depolarization are also observed in the *P*_1_, chaotic, and QP regimes. RGC model II displays akin results, SM Fig. S16(f). For STN in Fig. 10(b), SS occurs across a large portion of the parameter space (bellow the white dashed line in Fig. 9). But depolarization is restricted only to a portion of this region, gradually shifting to hyperpolarization as *A* increases.

**FIG. 10.**
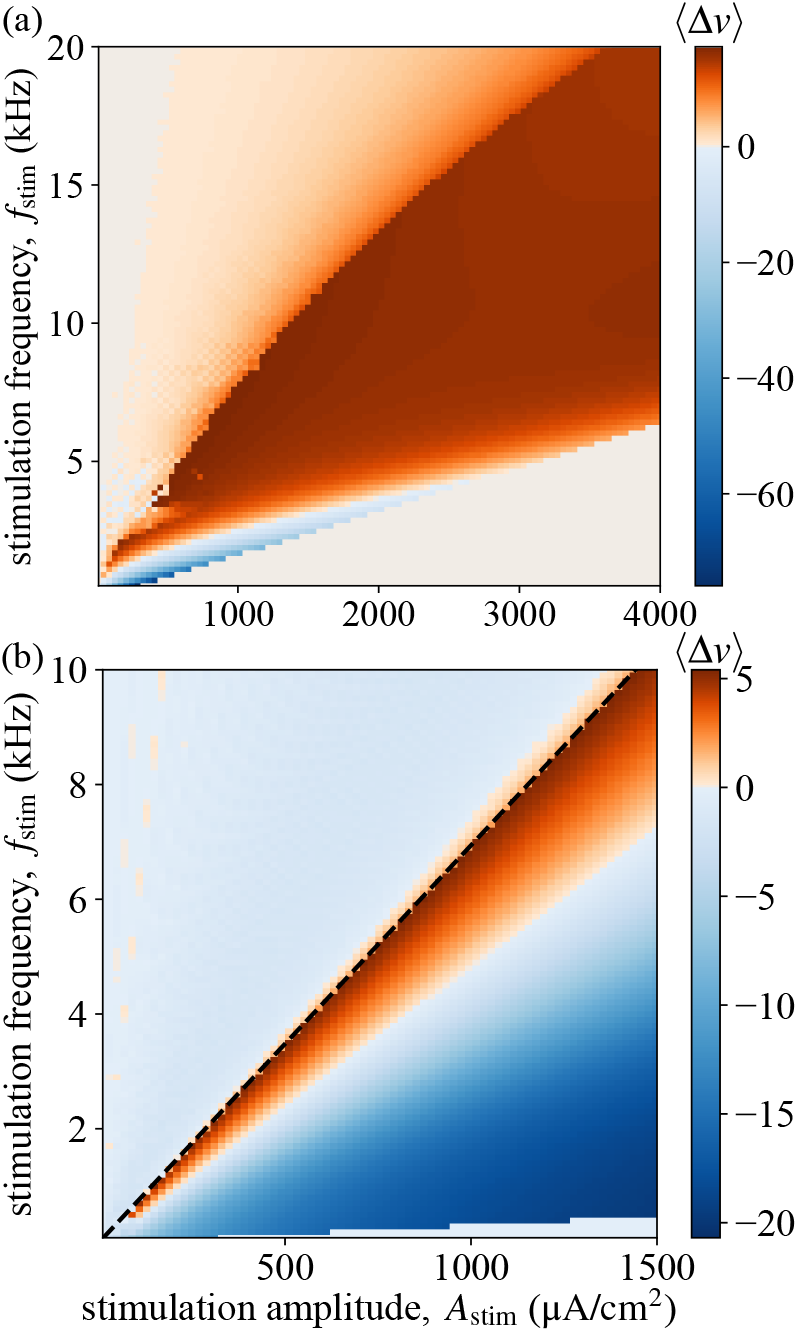
Parameter space mapping of the average membrane potential shift ⟨Δ*v*⟩ (mV) for mammalian neuronal models of (a) RGCs (model I) and (b) STN. Black dashed line in (b) marks the same value of *A* = 23 mV as the white dashed line in the STN functional map of Fig. 9. It divides the chaotic/QP and SS regions.

In HH, Cortical, Pyramidal Hippocampal and GPe models (SM Figs S16(a)-(c),(e)), depolarization mostly takes place in regions of high spiking activity, such as sub-harmonics, QP and chaotic. In contrast, basket hippocampal model (Fig. S16(d) in SM) displays stronger depolarization within SS regions, for *A* > 15.1 mV. These results indicate that while depolarization may contribute to SS in some neural cells, it is not a universal mechanism. We should emphasize that in [15], membrane potential dynamics were analyzed only for *I*_0_ = 0 and after the filtering procedure. Our functional mappings (Figs. 6 and 9) consider raw, unfiltered, responses.

### B. Physiological reliability of supra-physiological dynamical atlas

Conductance-based neuronal models are essentially built on physiological frequency 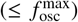 data. Thence, a natural question is if they are likewise able to replicate the dynamics in supra-physiological regimes. A thorough empirical study of the CNS under supra-physiological stimulation allied to comparisons with pertinent simulations, like those in the mappings and responses in Figs. 6 and 9, would be a way to settle the query. But such detailed investigations are absent in the literature. Thus, below we briefly review the current situation and remark on the physiological relevance of the present type of modeling.

There appears to be widespread consensus that conductance-based models combined with the cable equation may correctly replicate and forecast neuronal dynamics of UHF-stimulated PNS [23, 24, 41]. Nevertheless, as far as we know, no prior survey for CNS (an experimental challenge) has validated membrane potential dynamics under kilohertz considering conductance-based models. For instance, on the one hand, while data for hippocampal and cortical responses are available [16], theoretical modeling is lacking. On the other hand, STN neuronal model has been explored for kilohertz [17], but without experimental corroboration.

Given the previously mentioned scenario of insufficiency of theoretical-empirical comparisons under UHF stimulation, the fairly successful agreement of our results with a comprehensive modeling-data analyzes for retina neurons in [15] — refer to Sec. IV A — although a single example, already provides support for the physiological reliability of our findings. The fundamental point, however, is that even if the present models might not be able to accurately anticipate the precise locations and exact forms of the various dynamical regions in the stimulation parameter space (when faced with realistic experimental settings), they still can be of qualitative significance. Indeed, they provide a guide for the expected variety of dynamical responses and how they distribute across stimulation parameter space, helping to plan future experiments to validate CNS conductance models in the UHF regime.

As a final comment, we observe that in some models, e.g., standard HH, cortex, and hippocampus, the transition interfaces between distinct behaviors are not defined, displaying blending and mixing frontiers. Hence, these boundaries might be fractal, a recurrent phenomenon in various nonlinear dynamical systems [58]. But surely, when dealing with experimental data, these boundaries might not be obtained by simply considering pure dynamical systems methods, as deterministic rules are not available. Instead, one may need to introduce information-theoretical methods to treat such raw data, e.g., seeking to distinguish periodic and disordered (eventually noisy) responses [59, 60]. Also, from a modeling point of view, given the fast time scales associated with UHF, other temporal variations as stochastic ion channel fluctuations [61], should be taken into account.

### C. Technical aspects and scope of point neuron models

When modeling neuronal structures, two technical aspects are essential: the mathematical construction itself, which ideally should represent as realistically as possible the main mechanisms regulating the neuron’s actual functions and responses (see also the previous section); and the methods and simplifications assumed to solve the models (often involved, given their inherent nonlinearities). Our contribution points to few relevant issues along these lines.

Here, we have relied on biophysiologically motivated modeling, with a detailed description of ionic traits. Still, these models reduce the cell geometry to a single-compartment structure (a point). This reduction, although neglecting effects of axonal propagation [42] and external field coupling [62], has correctly predicted neuronal responses under kilohertz [15]. We observe that in contrast to multi-compartment, single-compartment neurons do not display the diffusion term ∂^2^*v*/∂*x*^2^ (for *x* the fiber axial direction). Thus, our response analysis is aimed to instances where the contributions of ionic currents and external stimulation are stronger than any local diffusion of the impulse. Such a regime is clearly appropriate to model patch-clamped setups [63]. However, the exact circumstances under which it occurs in external stimulation setups remain unclear (but refer to section IV D).

The non-linearity and high dimensionality of conductance-based models impose significant computational demands. The inclusion of time-dependent stimulation adds further complexity, making the system non-autonomous, most likely hindering analytical methods and solutions. In this regard, we mention averaging schemes to study non-autonomous systems [55]. In particular, they can be applied to conductance-based neuronal models when two conditions are met [17]: the neuron intrinsic spiking frequency under DC stimulation (*f*_osc_) is much lower than the stimulation frequency *f*_stim_; and (ii) the amplitude of induced membrane potential *A* remains small compared to typical spike amplitudes (*A* ≪ 100 mV). In our context, (i) is always true, as we work with supra-physiological frequencies. As for (ii), it is satisfied at low stimulation amplitudes, but might be violated at higher values. For instance, in Fig. 3(a) both conditions hold, once for (i) *f*_osc_ = 68.3 Hz ≪ *f*_stim_ = 1 kHz and for (ii) *A* = 11.9 mV ≪ 100 mV, so a regime well fitted for averaging techniques [17]. In contrast, in Fig. 3(d), where *A* ~ 95.5 mV, is not verified and replacing the original by an averaged system fails. These observations show that although averaging is useful for revealing some of the dynamics seen under kilohertz stimulation, great caution is required to ensure that such procedures comply with the parameters ranges investigated.

Hence, the dynamical systems methodologies employed in our work may be more appropriate for exploring the stimulation parameter space of increasingly intricate CNS neuron morphologies.

Additionally, dimensionality reduction or spike-resetting protocols are commonly used to mitigate computational complexity. Our findings show that in many instances to reduce the model dimensions significantly alters the dynamics in the supra-physiological stimulation regime, e.g., see Fig. 7(a)– (c). Moreover, albeit the influence of UHF stimulation on CNS network-level dynamics has not been addressed here, planned continuations of our study, either seeking to further characterize (a) the response of morphological CNS neurons to kilohertz, or trying to explain (b) empirical evidences [12] of therapeutic effects of kilohertz stimulation, should avoid reductions of fast gating variables. In particular, models based on reset conditions must be avoided near stimulated population, as they fail to represent the rapid membrane dynamics essential to (a) and (b).

### D. Similarities between the present dynamics and UHF stimulation of axons

As previously emphasized, axonal and dendritic components have been disregarded. Therefore, next we shall briefly compare the dynamics we have observed with already known results of PNS axons under UHF.

UHF stimulation of long axonal structures in the PNS elicits two main processes [42]. (1) Propagation block: axonal signal propagation is blocked near the stimulating electrode, where the stimulation current exceeds a threshold *I*_block_ [64]. The minimum frequency reported for this blocking is around 1 kHz [26, 65], but with no consensus on a limiting frequency, with proposed values of up to 100 kHz [10]. (2) Onset response: a brief asynchronous and bi-directional activity is triggered when the current exceeds *I*_onset_ (usually ≤ *I*_block_). This has been observed in animal experiments [23, 66] and *in-silico* models [24]. For motor neurons, the onset response involves an initial intense activity causing muscle twitch, followed by repetitive asynchronous activity, resulting in tetanic contractions. Although (2) is often viewed as a limitation of UHF stimulation, it has already been used to generate controllable, biomimetic responses [67].

Parallels to (1) and (2) above can be drawn for our CNS models. We observe large UHF-induced zones of spike suppression (either NS or SS), which somehow resemble propagation block, in the sense that external DC inputs fail to induce spikes. This phenomenon appears in six of the eight models investigated, the exceptions are GPe and pyramidal hippocampal. Besides, zones of aperiodic spiking activity (chaotic and QP) are observed, similarly to the onset response for axons. We hypothesize that similarly to [67], these responses could be used to control neural activity. Nevertheless, whether they work in an excitatory or inhibitory (related to desynchronization) manner is still up for debate. Finally, the emergence of subharmonic periodic regimes, *P*_*k*_, is rather surprising once no clear counterpart exists in axonal UHF dynamics. Interestingly, such behavior is well known in forced nonlinear oscillators [68].

## V. FINAL REMARKS AND CONCLUSION

In this contribution, we have considered a general framework based on dynamical systems to characterize how RGCs and five other CNS models respond to kilohertz forcing. Being a first effort towards this general goal, we have supposed structureless neurons, not taking into account certain morphological aspects. This obviously delimits the scope of our findings, clearly discussed in Secs. IV B and IV C. Through comprehensive simulations in a broad parameter space we have generated a systematic classification of the diverse set of dynamical responses of conductance-based neuronal models under rapid periodic stimulation — typical results are exemplified in the maps in Figs. 6 and 9. For example, we have described dynamical regimes related to kilohertz stimulation phenomena in RGC neurons, Sec. IV A, and PNS axons, Sec. IV D. Importantly, we have found that these responses are observed in different models of CNS neurons, providing an unifying comparison across mammalian cell models. We have also demonstrated that common reductions of sodium dynamics fail to describe phenomena under the kHz regime.

From our results, a set of issues arises, requesting further investigations so to provide a solid characterization of the UHF mechanism on CNS neurons. The most evident can be listed as the following. (a) To determine how the set of observed responses, described in our dynamical atlas, occur in multi-compartmental axonal structures under kilohertz extracellular stimulation. (b) To establish how these dynamical responses propagate, and how they are transmitted at the synapse. (c) To settle how different neural elements and morphologies — i.e., including soma, dendritic, and axonal structure — respond to external kHz stimulation. (d) Lastly, to understand how UHF regime of stimulation affects multiple scales of CNS structures.

Certainly, much work is necessary to elucidate (a)-(d) above, but associated to them we can mention potential suprathreshold neural effects to UHF responses (refer, e.g., to [10]). For instance, if the spike-suppression mechanism indeed blocks action potential propagation, it could block certain synaptic routes, altering network connectivity. Moreover, desynchronizing chaotic and quasi-periodic dynamics could influence excessive synchronization conditions. Finally, even if periodic sub-harmonic effects are not clearly understood, depending on their propagation properties, they still can potentially provide a different mechanism of neural excitation.

In conclusion, we hope to have established conductance-based models as the minimal, but relevant, theoretical construction for improving our comprehension of how CNS neurons respond to kilohertz stimulation.

## ACKNOWLEDGMENTS

First we would like to thank the anonymous referee for the rather useful suggestions, helping us to improve the quality of the discussions in the present contribution.

This study was financed in part by the Coordenação de Aperfeiçoamento de Pessoal de Nível Superior-Brazil (CAPES) - Finance Code 001 (Grant No. 88887.837508/2023-00) and from the project “Efficiency in uptake, production and distribution of photovoltaic energy distribution as well as other sources of renewable energy sources” (Grant No. 88881.311780/2018-00) via CAPES PRINT-UFPR. JGP acknowledges CAPES for a PhD scholarship (Grant No. 8887.927924/2023-00). MGELuz acknowledges CNPq-Brazil for the Research Grants No. 307512/2023-1 and No. 404577/2021-0 (Projeto Universal).

## Appendix A: Models

### 1. Hodgkin-Huxley model and dimension reduction

The Hodgkin-Huxley model [22] follows the electrical scheme presented in Fig. 1(b), where the membrane potential *v* is governed by

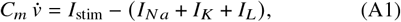

with *C*_*m*_ = 1 *μ*F · cm^−2^ the membrane capacitance per unit area. Moreover, *I*_*N a*_, *I*_*K*_ and *I*_*L*_, respectively, the sodium, potassium, and leakage, currents, read

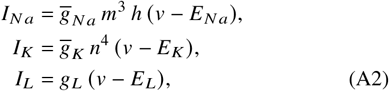

where 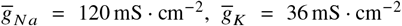, *g*_*L*_ = 0.3 mS · cm^−2^, *E*_*L*_ = −54.387 mV, *E*_*N a*_ = 50 mV and *E*_*K*_ = −77 mV.

The state of the HH model is given by **x** = [*v, m, h, n*]^*T*^, where *m, h*, ∈ [*n* 0, 1] are gating variables governing ion channel activation. For example, the value 1 (0) indicates fully open (closed) channel. Specifically, *m* and *h* control, respectively, the opening and closing of sodium channels, while *n* controls the opening of potassium channel.

The gating dynamics follows

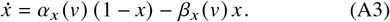

Here, *x* is either *m, h* or *n* and *α*_*x*_, *β*_*x*_ are voltage-dependent transition rates derived from experimental data in [22]. In this way, we have

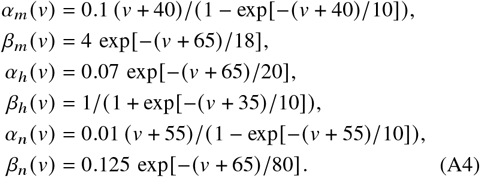

From Eqs. (A3) and (A4), we define the dimensionless steady-state *x*_∞_ and the time constant *τ*_*x*_ (in ms) as

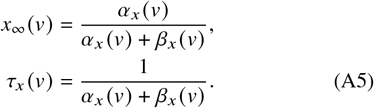

so that Eq. (A3) can be rewritten as

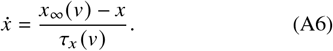

A common simplification results from *τ*_*m*_ ≪ *τ*_*h*_ and *τ*_*m*_ ≪ *τ*_*n*_. In such a regime, the system is reduced to three dimensions (3D), **x** = [*v, h, n*]^*T*^, by rewriting the sodium current as

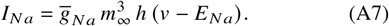

It is called the 3D *m* reduced model.

Another 3D reduction maintains the *m* dynamics, but considers a linear relation between *h* and *n*. Then, the state vector becomes **x** = [*v, m, n*]^*T*^, with the sodium current rewritten as

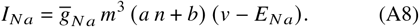

Numerically, *a* = − 1.073 and *b* = 0.877 (refer, e.g., to [38, 39]). For above one employs a least squares algorithm for the linear regression of *h* as function of *n*, assuming a constant *I*_stim_.

A further 2D reduction is possible by implementing two simplifications, namely, to set *m* = *m*_∞_ and to assume *h* = *an* + *b*, leading to **x** = [*v,n*] and a Na current in the form

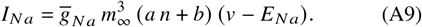

### 2. Models for the mammalian central nervous system

To explore the influence of UHF in a variety of locations in the mammalian central nervous system, we have considered mammalian brain models for the following regions: Basal Ganglia [46], modeling the Subthalamic Nucleus (STN) and Globus Pallidus externus (GPe); Cortex [43]; Hippocampal pyramidal excitatory [44] and basket inhibitory [45]; and finally Retinal Ganglion Cells (RGCs) [47].

In all cases, the membrane voltage *v* obeys an expression similar to Eq. (A1), or

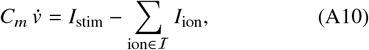

where ℐ is the set of channels, either ionic or leakage, and *I*_ion_ the associated current. The gating variables will depend on each specific central nervous system region. In Tab. I a summary of the models is given. It presents the dimension and state vector, channels current *I*_ion_’s equations, their conductance, reverse potential parameters, and the rates needed to describe the dynamics of auxiliary gating variables.

Almost all (see exceptions in SM Section S8) ionic currents *I*_ion_ have the form

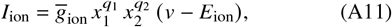

where 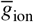 is the membrane ionic specific conductivity in mS · cm^−2^, *x*_1_ and *x*_2_ are gating variables, *q*_1_, *q*_2_ ≥ 0 are parameters defined for each model, and *E*_ion_ is the ion reversal potential. The gating variable *x*_*i*_ follows a differential equation akin to Eq. (A3). Likewise, *x*_∞_ (*v*) and *τ*_*x*_ (*v*) can be defined as in Eq. (A5), so that

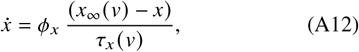

with *ϕ*_*x*_ an adjustment constant, shown on Table I only when *ϕ*_*x*_ ≠ 1.

Similarly to the original Hodgkin-Huxley model, a gating variable can be simplified, substituted by its steady-state value *x*_∞_. Such a process reduces the model dimensionality and is useful in physiological frequency regimes, but, as shown in Sec. III B, reductions must be performed rather carefully in the supra-physiological frequency regime. Important to emphasize that the mammalian models considered in our work are the same ones formulated and analyzed elsewhere. More concretely: for STN and GPe, as in [46] we set *m* = *m*_∞_, *a* = *a*_∞_, *b* = *b*_∞_, and *s* = *s*_∞_, which still leads to 5-dimensional systems; for Hippocampal inhibitory, we follow [45] and set *m* = *m*_∞_, reducing the model to 3D; lastly for Cortical [43], Hippocampal excitatory [44] and RGCs [47], no reductions are implemented. Some important governing equations are not exhausted by the relations in Table I. Hence, some extra details are given in SM Section S8.

Akin to the Hodgkin-Huxley, the mammalian models here can be classified as being either of type I or II [52], according to the DC current bifurcation analysis: the former exhibits a succession of saddle node points (or saddle-node homoclinic bifurcation) and can fire at arbitrarily low frequencies; whereas the latter exhibits a succession of Hopf bifurcations, firing only at a positive minimal frequency. The Supplementary Material section S5 shows the results of the bifurcation analysis for all the present models.

## Appendix B: Methods and Numerical Details

For the differential equations integration we have employed either the Python *scipy*-library [69] or the Julia Dynamical-System module [70]. Bifurcation analysis has been performed using XPPAUT 8.0. The codes used for simulations and figure generation are open-source and freely available at the GitHub repository [71].

### 1. Power spectral density

The power spectral density (PSD) of a discrete time series *X* = {*X*_0_, *X*_1_, …, *X*_*N* −1_}, with sampling period Δ*t* and time duration *T* = (*N* − 1) Δ*t*, measures the power density at each frequency band of *X*. It is defined as

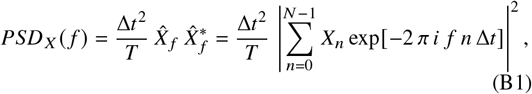

for 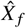 the Fourier transform of *X*. We have calculated 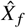 by the Fast Fourier Transform (FFT) method [72], as implemented in the NumPy library [73]. For the membrane voltage time series *v* (*t*), PSD is given in mV^2^ · Hz^−1^.

### 2. Lyapunov exponent

The Lyapunov exponent is a fundamental quantifier to typify chaotic trajectories. A trajectory is considered chaotic if it [74]: (a) is bounded, (b) is neither periodic nor consists of equilibrium points, and (c) exhibits at least one positive Lyapunov exponent. Conditions (a) and (b) can be qualitatively identified from trajectories and bifurcation diagrams, whereas condition (c) requires measuring at least one positive exponent for the trajectory. Therefore, to calculate the maximum Lyapunov exponent (MLE) suffices.

Two methods were employed to obtain the MLE of our systems. One is computationally fast and relies on averaging distances of nearby trajectories, obtaining only the MLE. The other, computationally expensive, calculates the full spectrum using the Jacobian of the system and QR-decomposition of perturbations. We provide detailed derivation of such methods in SM Section S9.

For the 4D HH and its lower-dimensional versions, we have employed the faster method. However, for mammalian models, we have found discrepancies between the two approaches for certain ranges of parameters. Hence, for mammalian models, the QR-method has been used. Both have been implemented in the `ChaosTools.jl` module of the `DynamicalSystems.jl` package [75] from Julia language [76].

Along the main text, we refer to the MLE simply as *λ*, independent on the protocol considered. Details of the simulations setups are found in the repository [71].

### 3. Membrane potential averaging method

To obtain the membrane potential average ⟨*v*⟩, we have adapted the method of [15]. First, we filter the external forcing artifact, then the entire filtered time series is averaged. The protocol starts by fitting a sinusoidal function *A* sin [2 *π f*_stim_ *t*+ *ϕ*] + *A*_0_ to *v* (*t*). Here, *f*_stim_ and *A* = *A*_stim_/ (2 *π f*_stim_) are fixed parameters, whereas the phase *ϕ* and intercept *A*_0_ are adjusted to *v t* by least squares method. Then, the filtered time series results from *v*_filtered_ *t* = *v t* − *A* sin [2 *π f*_stim_ *t + ϕ*] for *ϕ* the fitted phase. Finally, the time series of *v*_filtered_ is divided into *N* time steps of size Δ*t*, so ⟨*v*⟩ = 1/(*N* Δ*t*) _*n*_ *v*_filtered_(*n* Δ*t*).

### 4. Algorithm for behavioral mapping

To categorize the trajectories in terms of their possible dynamics, i.e., non-spiking (NS), spike-suppression (SS), chaotic, quasi-periodic (QP) and harmonics (*P*_*k*_), we have proceed as the following. First, the MLE *λ* is computed for a non-transient orbit and if *λ* > *λ*_min_, for *λ*_min_ a small threshold (here set to 10^−2^), the orbit is classified as chaotic. In the case of 0 < *λ* < *λ*_min_, the orbit is assumed quasi-periodic. If *λ* < 0, a trajectory is sampled at 1/*f*_stim_ time steps, leading to a stroboscopic orbit. Then, this orbit is tested for distinct periods *P*_*k*_: for a stroboscopic orbit *V*_strob_ = {*v*_1_, *v*_2_, … *v* _*N*_}, with *v*_*i*_ = *v* (*t* = *i*/*f*_stim_), we find the orbit to be of period *k* (so *P*_*k*_) for *k* the smallest integer such that |*v*_*n*_ − *v*_*n*+*k*_ | < *ϵ* for any *n* ≤ *N* − *k*. In particular, for *k* = 1 we have checked for the presence of spikes, using the sodium channel conductance similarly to the scheme used for ISI. If the orbit is *P*_1_, but there is no spike, we classify it either as NS or SS, depending on DC current behavior, otherwise it is considered *P*_1_.

## Notes

https://github.com/Jaderpolli/conductance-based-kHz

### Competing Interest Statement

The authors have declared no competing interest.

### Summary of Updates

Major modifications have been made to the Introduction section, avoiding previously excessive clinical contextualization, helping to focus and frame the manuscript as a contribution to forced dynamical systems under rapid forcing applied to neuronal dynamics. The discussion section has also been rearranged to focus on debating biophysiologically the results without direct clinical impacts. The quality of figures has been improved. Minor modifications in text and a few details from models in Appendix were moved to Supplemental Material.

## Reference

[1] C. W. Lynn and D. S. Bassett, The physics of brain network structure, function and control, Nature Reviews Physics 1, 318 (2019).

[2] P. Fries, Rhythms for cognition: communication through coherence, Neuron 88, 220 (2015).

[3] M. Thiebaut de Schotten and S. J. Forkel, The emergent properties of the connected brain, Science 378, 505 (2022).

[4] A. Fornito, A. Zalesky, and M. Breakspear, The connectomics of brain disorders, Nature Reviews Neuroscience 16, 159 (2015).

[5] E. S. Krames, P. H. Peckham, A. Rezai, and F. Aboelsaad, What is neuromodulation?, in Neuromodulation (Elsevier, 2009) pp. 3–8.

[6] A. D. Lawrence, A. H. Evans, and A. J. Lees, Compulsive use of dopamine replacement therapy in parkinson’s disease: reward systems gone awry?, The Lancet Neurology 2, 595 (2003).

[7] R. Rokyta and J. Fricová, Neurostimulation methods in the treatment of chronic pain, Physiological research 61, S23 (2012).

[8] J. D. Weiland and M. S. Humayun, Visual prosthesis, Proceedings of the IEEE 96, 1076 (2008).

[9] S. J. Bensmaia, D. J. Tyler, and S. Micera, Restoration of sensory information via bionic hands, Nature Biomedical Engineering 7, 443 (2023).

[10] C. Neudorfer, C. T. Chow, A. Boutet, A. Loh, J. Germann, G. J. Elias, W. D. Hutchison, and A. M. Lozano, Kilohertz-frequency stimulation of the nervous system: A review of underlying mechanisms, Brain Stimulation: Basic, Translational, and Clinical Research in Neuromodulation 14, 513 (2021).

[11] A. Soin, Z.-P. Fang, and J. Velasco, Peripheral neuromodulation to treat postamputation pain, Stimulation of the Peripheral Nervous System 29, 158 (2016).

[12] I. E. Harmsen, D. J. Lee, R. F. Dallapiazza, P. De Vloo, R. Chen, A. Fasano, S. K. Kalia, M. Hodaie, and A. M. Lozano, Ultrahigh-frequency deep brain stimulation at 10,000 Hz improves motor function, Movement Disorders 34, 146 (2019).

[13] S. V. Karnup, W. De Groat, J. Beckel, and C. Tai, Effect of biphasic khz field stimulation on ca1 pyramidal neurons in slices, Medical Research Archives 10, 10.18103/mra.v10i8.2957 (2022).

[14] L. S. Lesperance, M. Lankarany, T. C. Zhang, R. Esteller, S. Ratté, and S. A. Prescott, Artifactual hyperpolarization during extracellular electrical stimulation: Proposed mechanism of high-rate neuromodulation disproved, Brain stimulation 11, 582 (2018).

[15] J.-I. Lee, P. Werginz, T. Kameneva, M. Im, and S. I. Fried, Membrane depolarization mediates both the inhibition of neural activity and cell-type-differences in response to high-frequency stimulation, Communications Biology 7, 734 (2024).

[16] C. R. Ravasio, K. Kondabolu, S. Zhou, E. Lowet, E. San Antonio, R. A. Mount, S. K. Bhogal, and X. Han, Kilohertz electrical stimulation evokes robust cellular responses like conventional frequencies but distinct population dynamics, Communications Biology 8, 1 (2025).

[17] K. Pyragas, V. Novičenko, and P. A. Tass, Mechanism of suppression of sustained neuronal spiking under high-frequency stimulation, Biological Cybernetics 107 (2013).

[18] M. Popova, C. C. Hilgetag, and M.-T. Hütt, Perturbation therapies for neurodegenerative disorders: How attractors of excitable networks can help, Physical Review E 110, 054406 (2024).

[19] M. Lucas, J. Newman, and A. Stefanovska, Stabilization of dynamics of oscillatory systems by nonautonomous perturbation, Physical Review E 97, 042209 (2018).

[20] P. A. Tass, L. Qin, C. Hauptmann, S. Dovero, E. Bezard, T. Boraud, and W. G. Meissner, Coordinated reset has sustained aftereffects in parkinsonian monkeys, Annals of neurology 72, 816 (2012).

[21] J. Wang, S. Nebeck, A. Muralidharan, M. D. Johnson, J. L. Vitek, and K. B. Baker, Coordinated reset deep brain stimulation of subthalamic nucleus produces long-lasting, dosedependent motor improvements in the 1-methyl-4-phenyl-1, 2, 3, 6-tetrahydropyridine non-human primate model of parkinsonism, Brain stimulation 9, 609 (2016).

[22] A. L. Hodgkin and A. F. Huxley, A quantitative description of membrane current and its application to conduction and excitation in nerve, The Journal of physiology 117, 500 (1952).

[23] K. L. Kilgore and N. Bhadra, Nerve conduction block utilising high-frequency alternating current, Medical and Biological Engineering and Computing 42, 394 (2004).

[24] N. Bhadra, E. A. Lahowetz, S. T. Foldes, and K. L. Kilgore, Simulation of high-frequency sinusoidal electrical block of mammalian myelinated axons, Journal of computational neuroscience 22, 313 (2007).

[25] M. H. Negm and I. C. Bruce, The effects of hcn and klt ion channels on adaptation and refractoriness in a stochastic auditory nerve model, IEEE Transactions on Biomedical Engineering 61, 2749 (2014).

[26] N. Bhadra and K. L. Kilgore, High-frequency electrical conduction block of mammalian peripheral motor nerve, Muscle & Nerve: Official Journal of the American Association of Electrodiagnostic Medicine 32, 782 (2005).

[27] C. Tai, J. R. Roppolo, and W. C. de GROAT, Response of external urethral sphincter to high frequency biphasic electrical stimulation of pudendal nerve, The Journal of urology 174, 782 (2005).

[28] N. Bhadra, E. L. Foldes, D. M. Ackermann, and K. L. Kilgore, Reduction of the onset response in high frequency nerve block with amplitude ramps from non-zero amplitudes, in 2009 Annual International Conference of the IEEE Engineering in Medicine and Biology Society (IEEE, 2009) pp. 650–653.

[29] N. Pelot and W. Grill, In vivo quantification of excitation and kilohertz frequency block of the rat vagus nerve, Journal of neural engineering 17, 026005 (2020).

[30] B. Hassard, Bifurcation of periodic solutions of the hodgkinhuxley model for the squid giant axon, Journal of Theoretical Biology 71, 401 (1978).

[31] I. S. Labouriau, Degenerate hopf bifurcation and nerve impulse, SIAM journal on mathematical analysis 16, 1121 (1985).

[32] J. Guckenheimer and J. Labouriau, Bifurcation of the hodgkin and huxley equations: a new twist, Bulletin of Mathematical Biology 55, 937 (1993).

[33] H. Fukai, T. Nomura, S. Doi, and S. Sato, Hopf bifurcations in multiple-parameter space of the hodgkin-huxley equations ii. singularity theoretic approach and highly degenerate bifurcations, Biological Cybernetics 82, 223 (2000).

[34] K. Aihara, G. Matsumoto, and Y. Ikegaya, Periodic and nonperiodic responses of a periodically forced Hodgkin-Huxley oscillator, Journal of Theoretical Biology 109, 249 (1984).

[35] S.-G. Lee and S. Kim, Bifurcation analysis of mode-locking structure in a hodgkin-huxley neuron under sinusoidal current, Phys. Rev. E 73, 041924 (2006).

[36] L. S. Borkowski, Bistability and resonance in the periodically stimulated hodgkin-huxley model with noise, Phys. Rev. E 83, 051901 (2011).

[37] B. A. y Arcas, A. L. Fairhall, and W. Bialek, Computation in a single neuron: Hodgkin and huxley revisited, Neural computation 15, 1715 (2003).

[38] T. B. Kepler, L. Abbott, and E. Marder, Reduction of conductance-based neuron models, Biological cybernetics 66, 381 (1992).

[39] C. Meunier, Two and three dimensional reductions of the hodgkin-huxley system: separation of time scales and bifurcation schemes, Biological cybernetics 67, 461 (1992).

[40] B. Ermentrout, Reduction of conductance-based models with slow synapses to neural nets, Neural Computation 6, 679 (1994).

[41] D. M. Ackermann, N. Bhadra, M. Gerges, and P. J. Thomas, Dynamics and sensitivity analysis of high-frequency conduction block, Journal of neural engineering 8, 065007 (2011).

[42] Y. A. Patel and R. J. Butera, Challenges associated with nerve conduction block using kilohertz electrical stimulation, Journal of neural engineering 15, 031002 (2018).

[43] A. Erisir, D. Lau, B. Rudy, and C. S. Leonard, Function of specific k+ channels in sustained high-frequency firing of fastspiking neocortical interneurons, Journal of neurophysiology 82, 2476 (1999).

[44] R. D. Traub and R. Miles, Neuronal networks of the hippocampus, Vol. 777 (Cambridge University Press, 1991).

[45] X.-J. Wang and G. Buzsáki, Gamma oscillation by synaptic inhibition in a hippocampal interneuronal network model, Journal of neuroscience 16, 6402 (1996).

[46] D. Terman, J. E. Rubin, A. Yew, and C. Wilson, Activity patterns in a model for the subthalamopallidal network of the basal ganglia, Journal of Neuroscience 22, 2963 (2002).

[47] J. F. Fohlmeister, E. D. Cohen, and E. A. Newman, Mechanisms and distribution of ion channels in retinal ganglion cells: using temperature as an independent variable, Journal of neurophysiology 103, 1357 (2010).

[48] J. C. Brumberg and B. S. Gutkin, Cortical pyramidal cells as non-linear oscillators: experiment and spike-generation theory, Brain research 1171, 122 (2007).

[49] K. Tateno, H. Hayashi, and S. Ishizuka, Complexity of spatiotemporal activity of a neural network model which depends on the degree of synchronization, Neural Networks 11, 985 (1998).

[50] S. Ahn, S. E. Zauber, R. M. Worth, and L. L. Rubchinsky, Synchronized beta-band oscillations in a model of the globus pallidus-subthalamic nucleus network under external input, Frontiers in computational neuroscience 10, 134 (2016).

[51] D. W. Crevier and M. Meister, Synchronous period-doubling in flicker vision of salamander and man, Journal of neurophysiology 79, 1869 (1998).

[52] B. Ermentrout and D. H. Terman, Mathematical foundations of neuroscience, Vol. 35 (Springer, 2010).

[53] H. Sakaguchi and K. Yamasaki, Suppression and frequency control of repetitive spiking in the FitzHugh-Nagumo model, Physical Review E 108, 014207 (2023).

[54] S. H. Strogatz, Nonlinear dynamics and chaos: with applications to physics, biology, chemistry, and engineering (CRC press, 2018).

[55] J. A. Sanders, F. Verhulst, and J. Murdock, Averaging methods in nonlinear dynamical systems, Vol. 59 (Springer, 2007).

[56] T. Kameneva, M. Maturana, A. Hadjinicolaou, S. Cloherty, M. Ibbotson, D. Grayden, A. Burkitt, and H. Meffin, Retinal ganglion cells: mechanisms underlying depolarization block and differential responses to high frequency electrical stimulation of on and off cells, Journal of neural engineering 13, 016017 (2016).

[57] T. Guo, D. Tsai, C. Y. Yang, A. Al Abed, P. Twyford, S. I. Fried, J. W. Morley, G. J. Suaning, S. Dokos, and N. H. Lovell, Mediating retinal ganglion cell spike rates using high-frequency electrical stimulation, Frontiers in neuroscience 13, 413 (2019).

[58] J. Aguirre, R. L. Viana, and M. A. Sanjuán, Fractal structures in nonlinear dynamics, Reviews of Modern Physics 81, 333 (2009).

[59] H. Kantz, A robust method to estimate the maximal lyapunov exponent of a time series, Physics letters A 185, 77 (1994).

[60] J. V. V. Flauzino, T. L. Prado, N. Marwan, J. Kurths, and S. R. Lopes, Quantifying disorder in data, Physical Review Letters 135, 097401 (2025).

[61] J. A. White, J. T. Rubinstein, and A. R. Kay, Channel noise in neurons, Trends in neurosciences 23, 131 (2000).

[62] T. Caussade, E. Paduro, M. Courdurier, E. Cerpa, W. M. Grill, and L. E. Medina, Towards a more accurate quasi-static approximation of the electric potential for neurostimulation with kilohertz-frequency sources, Journal of neural engineering 20, 066035 (2023).

[63] C. Koch and I. Segev, Methods in neuronal modeling: from ions to networks (MIT press, 1998).

[64] K. L. Kilgore and N. Bhadra, Reversible nerve conduction block using kilohertz frequency alternating current, Neuromodulation: Technology at the Neural Interface 17, 242 (2014).

[65] R. P. Williamson and B. J. Andrews, Localized electrical nerve blocking, IEEE Transactions on Biomedical Engineering 52, 362 (2005).

[66] B. R. Bowman and D. R. McNeal, Response of single alpha motoneurons to high-frequency pulse trains: firing behavior and conduction block phenomenon, Stereotactic and Functional Neurosurgery 49, 121 (1986).

[67] E. Formento, E. D’Anna, S. Gribi, S. P. Lacour, and S. Micera, A biomimetic electrical stimulation strategy to induce asynchronous stochastic neural activity, Journal of neural engineering 17, 046019 (2020).

[68] C. Hayashi, Nonlinear oscillations in physical systems (Princeton University Press, 2014).

[69] P. Virtanen, R. Gommers, T. E. Oliphant, M. Haberland, T. Reddy, D. Cournapeau, E. Burovski, P. Peterson, W. Weckesser, J. Bright, S. J. van der Walt, M. Brett, J. Wilson, K. J. Millman, N. Mayorov, A. R. J. Nelson, E. Jones, R. Kern, E. Larson, C. J. Carey, İ. Polat, Y. Feng, E. W. Moore, J. VanderPlas, D. Laxalde, J. Perktold, R. Cimrman, I. Henriksen, E. A. Quintero, C. R. Harris, A. M. Archibald, A. H. Ribeiro, F. Pedregosa, P. van Mulbregt, and SciPy 1.0 Contributors, SciPy 1.0: Fundamental Algorithms for Scientific Computing in Python, Nature Methods 17, 261 (2020).

[70] G. Datseris, Dynamicalsystems.jl: A julia software library for chaos and nonlinear dynamics, Journal of Open Source Software 3, 598 (2018).

[71] J. G. Polli, F. Kolbl, M. E. E. da Luz, and P. Lanusse, Jaderpolli/conductance-based-khz: cb-khz (2025).

[72] J. W. Cooley and J. W. Tukey, An algorithm for the machine calculation of complex fourier series, Mathematics of computation 19, 297 (1965).

[73] C. R. Harris, K. J. Millman, S. J. van der Walt, R. Gommers, P. Virtanen, D. Cournapeau, E. Wieser, J. Taylor, S. Berg, N. J. Smith, R. Kern, M. Picus, S. Hoyer, M. H. van Kerkwijk, M. Brett, A. Haldane, J.F. delRío, M. Wiebe, P. Peterson, P. Gérard-Marchant, K. Sheppard, T. Reddy, W. Weckesser, H. Abbasi, C. Gohlke, and T. E. Oliphant, Array programming with NumPy, Nature 585, 357 (2020).

[74] K. T. Alligood, T. D. Sauer, J. A. Yorke, and D. Chillingworth, Chaos: an introduction to dynamical systems, SIAM Review 40, 732 (1998).

[75] G. Datseris, Dynamicalsystems. jl: A julia software library for chaos and nonlinear dynamics, Journal of Open Source Software 3, 598 (2018).

[76] J. Bezanson, A. Edelman, S. Karpinski, and V. B. Shah, Julia: A fresh approach to numerical computing, SIAM review 59, 65 (2017).

