## Supplementar Material for "Dynamical Diversity in Conductance-Based Neuron Response to kilohertz Electrical Stimulation"

### SUPPLEMENTARY TEXT

#### S1. Extra Stroboscopic Bifurcation Diagrams

In this section, additional stroboscopic bifurcation diagrams for different parameters of stimulation frequency $f_{\text{stim}}$ and constant input $I_0$ are provided for the HH model. Figure S1 compares stroboscopic diagrams when the system is stimulated with DC $I_0 = 0\,\mu\text{A/cm}^2$ and with a $I_0 = 10\,\mu\text{A/cm}^2$ at different values of supra-physiological frequencies. Namely, $f_{\text{stim}} = 1.0\,\text{kHz}$ in (a) and (b), $f_{\text{stim}} = 1.25\,\text{kHz}$ in (c) and (d), $f_{\text{stim}} = 1.65\,\text{kHz}$ in (e) and (f), and $f_{\text{stim}} = 2.75\,\text{kHz}$ in (g) and (h).

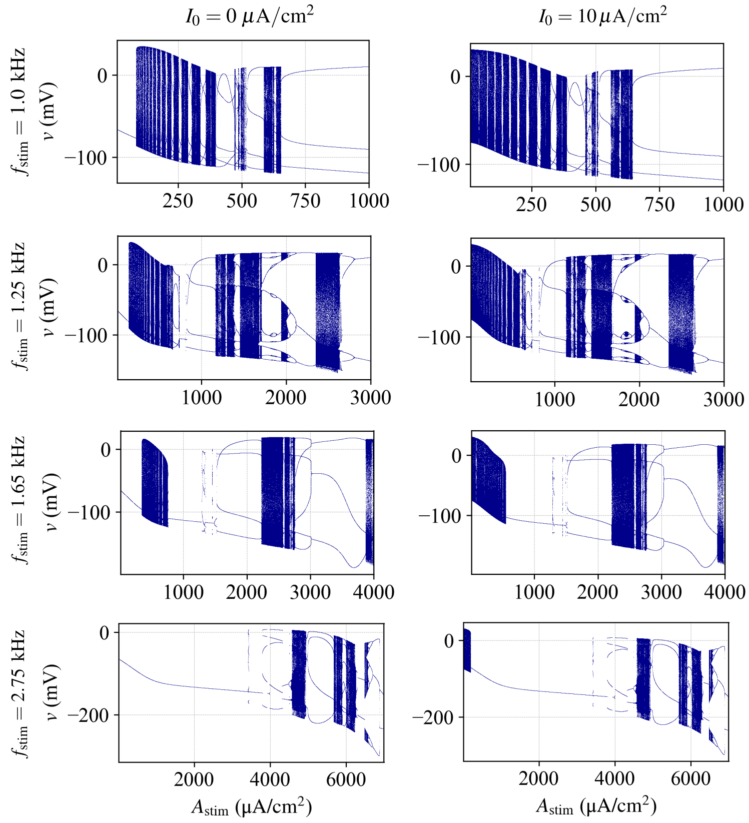


FIG. S1: Stroboscopic bifurcation diagrams of HH-model for additional parameters of frequency of stimulation $f_{\text{stim}}$ and constant stimulation input $I_0$. In (a), (c), (e), (g) the external constant input $I_0 = 0$ and, respectively, $f_{\text{stim}} = 1.0$, 1.25, 1.65, and 2.75 kHz. In (b), (d), (f), (h) the external constant input is $I_0 = 10\,\mu\text{A/cm}^2$ and, respectively $f_{\text{stim}} = 1.0$, 1.25, 1.65, and 2.75 kHz.

For all tested cases, when $I_0 = 0\,\mu\text{A/cm}^2$, no spike is observed for small values of $A_{\text{stim}}$, while for $I_0 = 10\,\mu\text{A/cm}^2$ even for $A_{\text{stim}} = 0.0\,\mu\text{A/cm}^2$ there are spiking dynamics. This is expected for the HH-model since, as the equilibrium point is stable for all $I_0 < I_{H_1} = 9.775357$, while for $I_0 > I_{H_1}$ up to $I_0 = I_{H_2} = 154.5226$ there is a stable limit cycle, i.e., spiking dynamics. Remarkably, as $A_{\text{stim}}$ increases further, the dynamics has almost no influence from the constant input, since the sinusoidal stimulation dominates the external forcing.

### S2. Bifurcation Analysis for reduced models with constant stimulation

In this section, we show the bifurcation diagrams, ISI, and spiking frequency for the reduced HH models. Figs. S2, S3 and S4 show the DC stimulation analysis of ISI, spiking/oscillation frequency, and bifurcation diagram for the 3D-$h$, 3D-$m$, and 2D-reduction models, respectively.

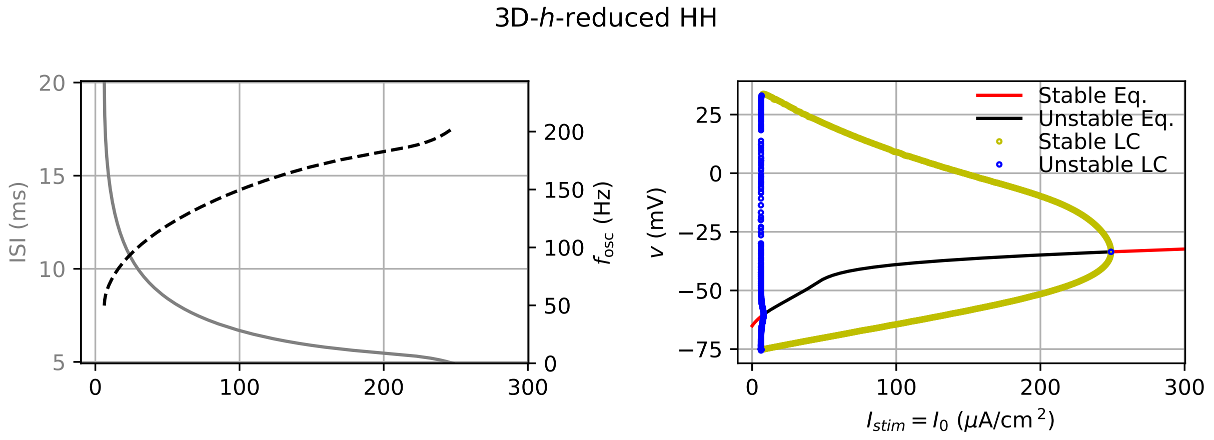


FIG. S2: Model properties of the 3D-$h$ reduced HH model. On the left, ISI and spiking frequency, on the right, bifurcation diagram.

All models show positive minimal spiking frequency after a subcritical hopf bifurcation at $I_0^{H_1}$, and the spiking behavior continues until the supercritical hopf bifurcation at $I_0^{H_2}$. The values of critical DC current $I_0^{H_1}$, $I_0^{H_2}$, and maximal frequency of oscillation $f_{\text{osc}}^{\max}$, for each model, including the original HH model, are display at Table S1

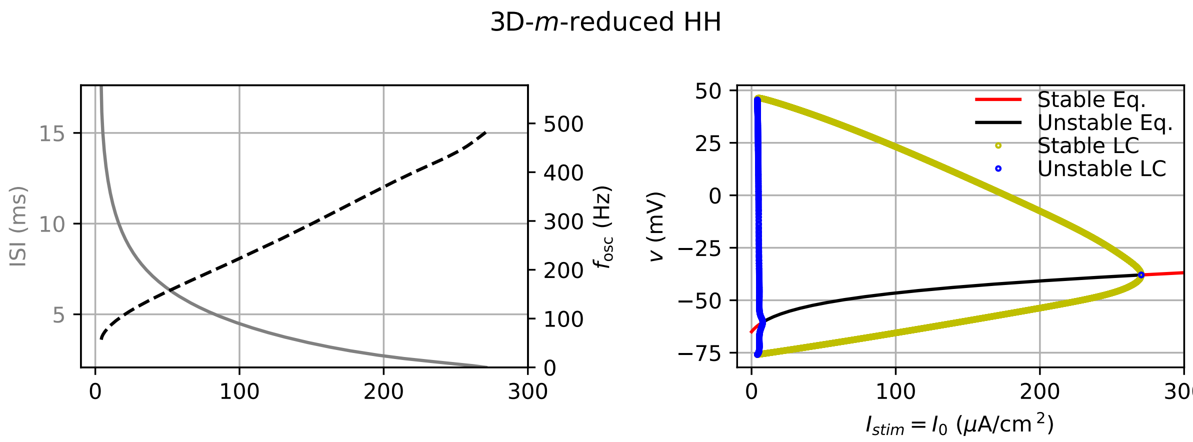


FIG. S3: Model properties of the 3D-$m$ reduced HH model. On the left, ISI and spiking frequency, on the right, bifurcation diagram.

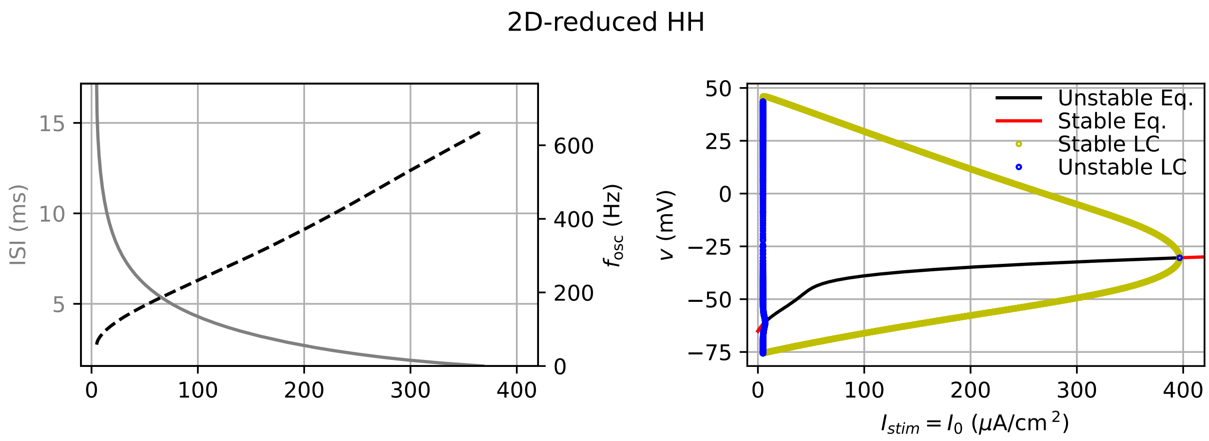


FIG. S4: Model properties of the 2D reduced HH model. On the left, ISI and spiking frequency, on the right, bifurcation diagram.

| Model | $I_0^{H1}$ ($\mu$A/cm$^2$) | $I_0^{H2}$ ($\mu$A/cm$^2$) | $f_{\text{osc}}^{\text{max}}$ (Hz) |
|---|---|---|---|
| 4D-HH | 9.775 | 154.5 | 169.0 |
| 3D-$h$ reduced HH | 7.966 | 249.0 | 202.9 |
| 3D-$m$ reduced HH | 7.743 | 270.3 | 481.8 |
| 2D reduced HH | 6.844 | 396.7 | 639.7 |

TABLE S1: Bifurcation points and maximum oscillatory frequencies of original Hodgkin Huxley model and various simplified models (all numbers rounded at 4 significant digits)

#### S3. Lyapunov Exponent for 3D-Reduced models

The Lyapunov Exponent of the 3D-$h$-reduced and 3D-$m$-reduced models is shown in Figs. S5 and S6, respectively. The 2D reduction is omitted as no regions of $\lambda > 0$ are observed.

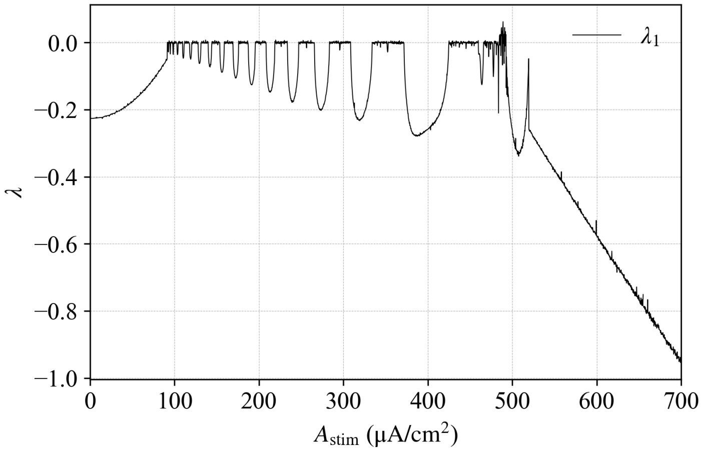


FIG. S5: Lyapunov for the 3D-$h$-reduced model.

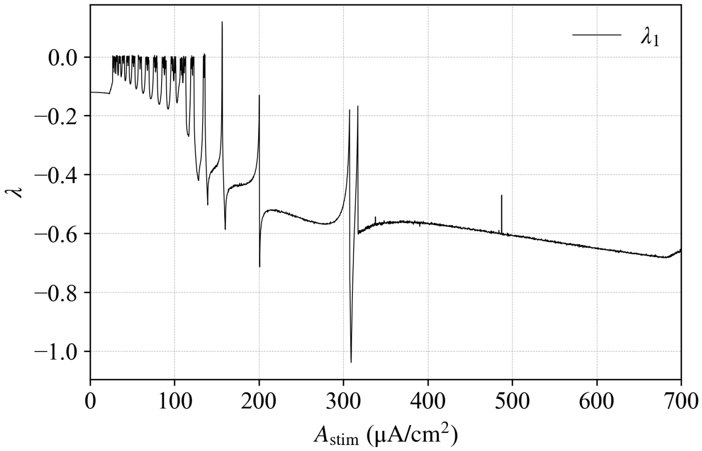


FIG. S6: Lyapunov for the 3D-$m$-reduced model.

### S4. Stroboscopic Bifurcation Diagram for 2D-reduced model

Fig. S7 shows the bifurcation diagram for the 2D-reduced HH model under stimulation frequency $f_{\text{stim}} = 1\,\text{kHz}$ and amplitude range $A_{\text{stim}} \in [0, 700]\,\mu\text{A/cm}^2$. The 2D-reduction considers $m = m_\infty$ and a linear relation between $h$ and $n$ gating variables $h = an + b$. An alternating sequence of quasiperiodic and periodic behaviors is observed for $A_{\text{stim}} \lesssim 100\,\mu\text{A/cm}^2$, closely matching the 3D-$m$-reduction and appearing earlier than in 3D-$h$-reduction and original 4D model. Since continuous-time dynamical systems with $d < 3$ cannot exhibit chaos, computing the MLE for this 3D-reduced model is unnecessary. The similarity between 2D with the 3D-$m$-reduction highlights the critical role of sodium activation dynamics — represented by the $m$ variable — in shaping the response under supra-physiological frequency inputs.

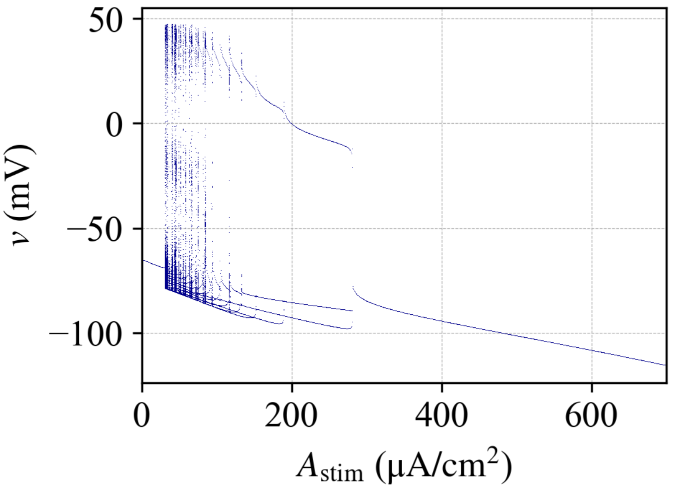


FIG. S7: Stroboscopic bifurcation diagram for the 2D-reduced HH-model when stimulated at $f_{\text{stim}} = 1\,\text{kHz}$ frequency and $A_{\text{stim}} = [0, 700]\,\mu\text{A/cm}^2$ amplitude range.

### S5. Model properties for constant stimulation of mammalian models

#### 1. *Type I*

A sequence of specific bifurcations gives rise to the behavior observed in type I models. First, a sequence of two saddle-node bifurcations on an invariant cycle (SNIC) occurs at DC values $I_0^{SN1}$ and $I_0^{SN2}$. Then, a stable limit cycle (LC), where spiking activity occurs, is maintained up to a hopf bifurcation point $I_0^H$. The stable LC is maintained for DC values $I_0^{SN1} \leq I_0 \leq I_0^H$.

We used various models exhibiting such behavior: Cortical (see fig. S8), pyramidal excitatory (see fig. S9), basket inhibitory (see fig. S10) and STN models (see fig. S11).

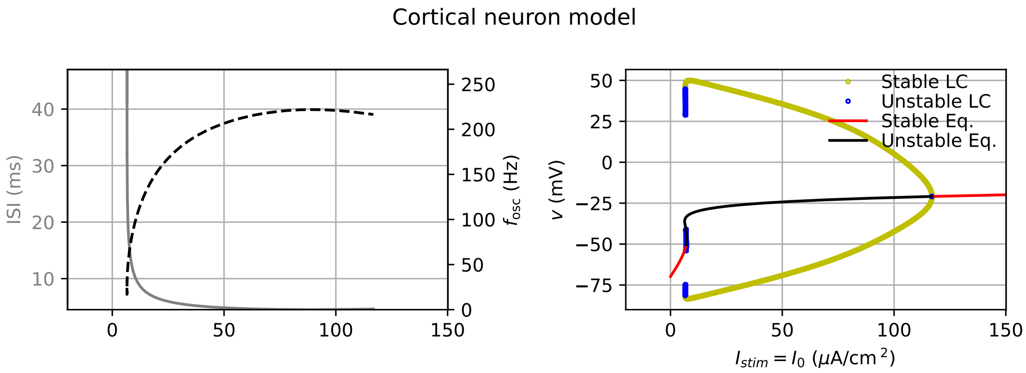


FIG. S8: Model properties of the cortical model. On the left, ISI and spiking frequency, on the right, bifurcation diagram.

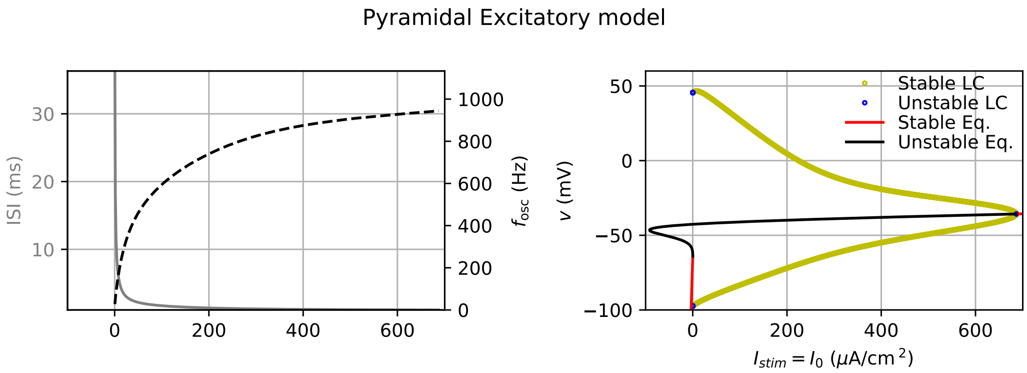


FIG. S9: Model properties of the piramidal excitatory model. On the left, ISI and spiking frequency, on the right, bifurcation diagram.

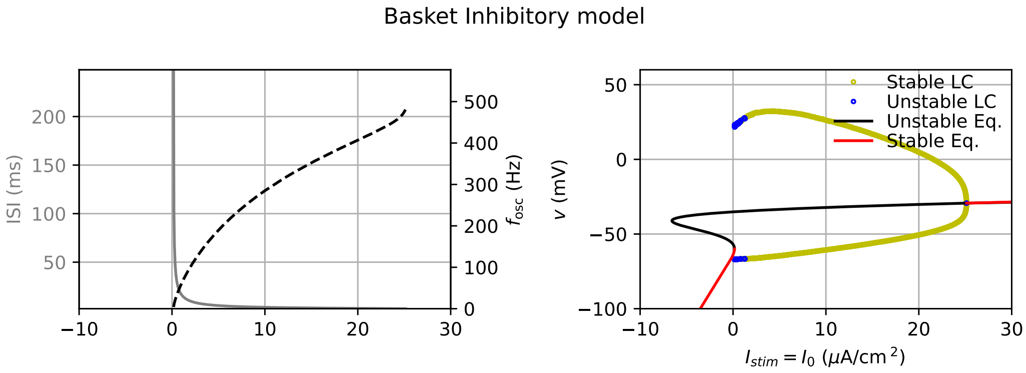


FIG. S10: Model properties of the basket inhibitory model. On the left, ISI and spiking frequency, on the right, bifurcation diagram.

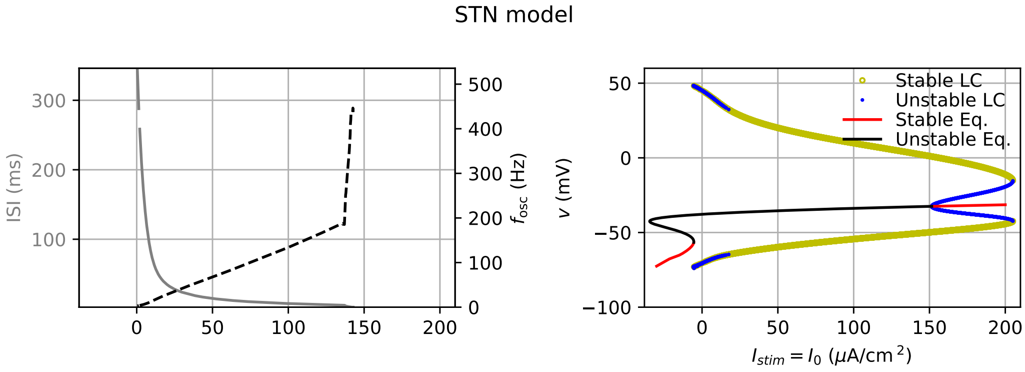


FIG. S11: Model properties of the STN model. On the left, ISI and spiking frequency, on the right, bifurcation diagram.

Values of stimulation and maximum oscillatory frequency are summed up in table S2.

| *model* | $I_0^{SN1}$ ($\mu$A.cm$^{-2}$) | $I_0^{SN2}$ ($\mu$A.cm$^{-2}$) | $I_0^{H}$ ($\mu$A.cm$^{-2}$) | $f_{\text{osc}}^{max}$ (Hz) |
|---|---|---|---|---|
| cortical | 6.304 | 7.406 | 116.9 | 221.9 |
| pyramidal excitatory | $-91.63$ | $1.193 \times 10^{-1}$ | 686.8 | 994.3 |
| basket inhibitory | $-6.579$ | $1.160 \times 10^{-1}$ | 25.13 | 480.7 |
| STN | $-34.59$ | $-5.465$ | 151.5 | 446.4 |

TABLE S2: Bifurcation points and maximum oscillatory frequencies of type I models (all numbers rounded at 4 significant digits).

*2. Type II*

Type II models are characterized by a succession of two Hopf bifurcation ($H1$ corresponding to $I_0^{H1}$ and $H2$ corresponding to $I_0^{H1}$) and a Hopf bifurcation ($H$ corresponding to $I_0^H$). The studied GPe model exhibits such behavior (see fig. S12). Values of stimulation and maximum oscillatory frequency are summed up in table S3.

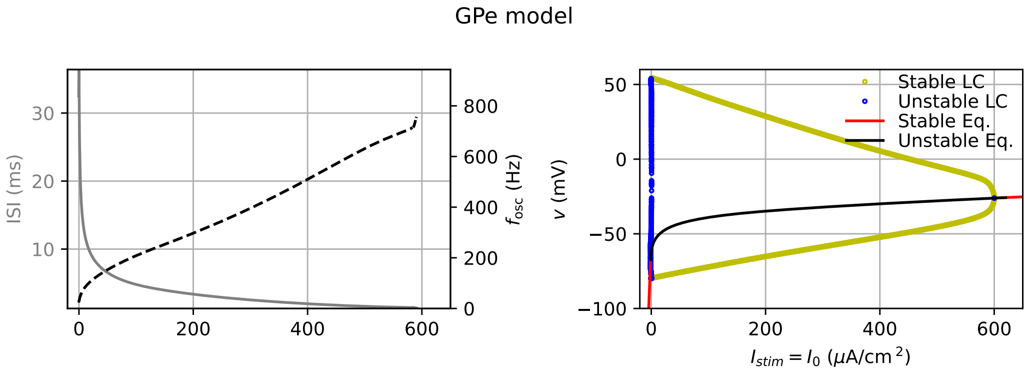


FIG. S12: Model properties of the GPe model - upper graph: ISI and spiking frequency, lower graph: bifurcation diagram

| *model* | $I_0^{H1}$ ($\mu$A.cm$^{-2}$) | $I_0^{H2}$ ($\mu$A.cm$^{-2}$) | $f_{\text{osc}}^{max}$ (Hz) |
|---|---|---|---|
| GPe | $-1.031$ | 600.2 | 786.1 |

TABLE S3: Bifurcation points and maximum oscillatory frequencies of type II models (all numbers rounded at 4 significant digits)

*3. Note on RGC 1 and 2 models*

The analysis of RGC models (1 and 2) was not successfull using the XPP. Authors could not systematically converge to eigenvalues on the range of stimulation current where a stable nor an oscillatory behavior was observed. We provide the ISI and oscillatory frequency curves in fig. S13 and the following figures: for RGC 1 model, the maximal oscillatory frequency is $f_{osc}^{max} = 1522$Hz and for RGC 2 model, the maximal oscillatory frequency is $f_{osc}^{max} = 1990$Hz

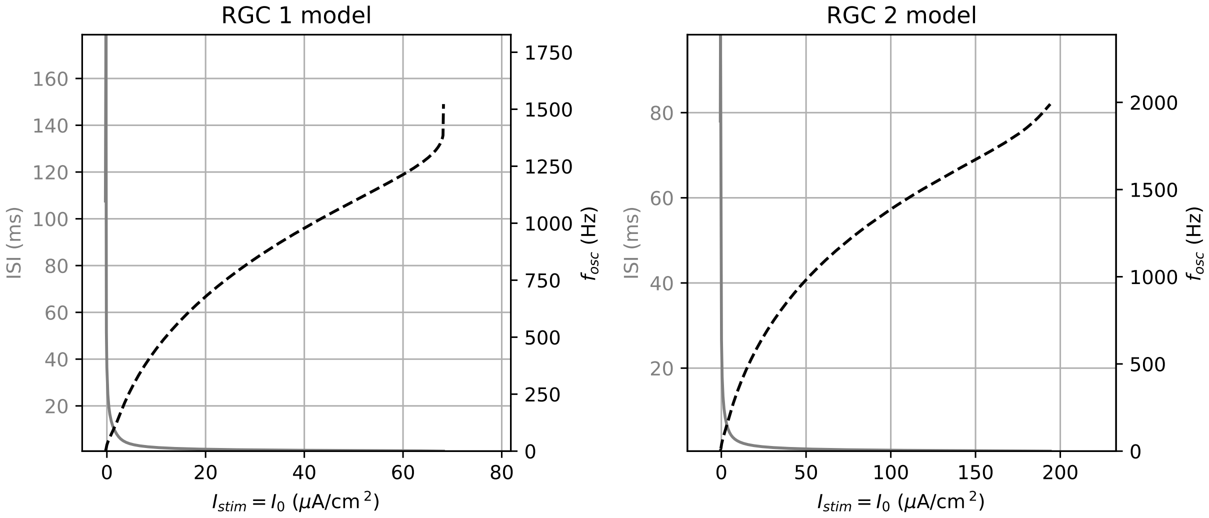


FIG. S13: ISI and spiking frequency for RGC 1 and 2 models

#### *4. Transition frequencies of particles*

All transition frequencies for the particles considered in the mammalian models used in the results are shown in Fig. S14.

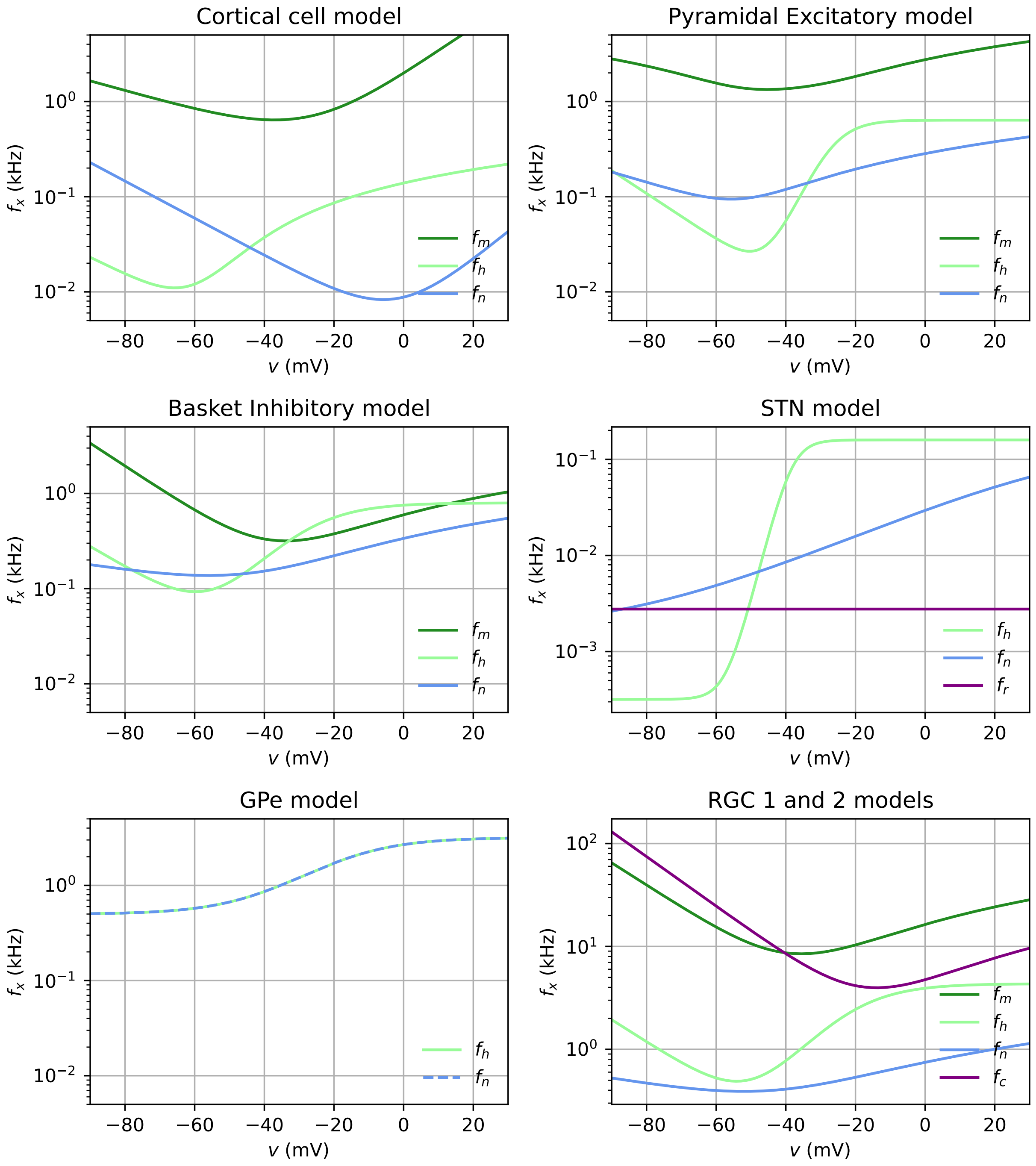


FIG. S14: Transition frequencies $f_x$ of gating variables for the mammalian CNS models.

### S6. Functional mapping for non-spiking inducing DC

For when the DC stimulation $I_0$ parameter does not elicit spikes alone, we can observe non-spiking (NS) dynamics in mammalian models, in the same way as observed in the HH model with $I_0 = 0$. Figure S15 shows the functional mapping for all mammalian models, in a parallel with Fig. 9 of main text. Here, we use values of $I_0 < I_0^{SN2}$ for type I models, and $I_0 < I_0^{H_1}$ for type II models. Specifically, for STN $I_0 = -10\,\mu\text{A/cm}^2$, for RGCs and GPe $I_0 = -5\,\mu\text{A/cm}^2$, while for Cortex and both Hippocampal models $I_0 = 0$.

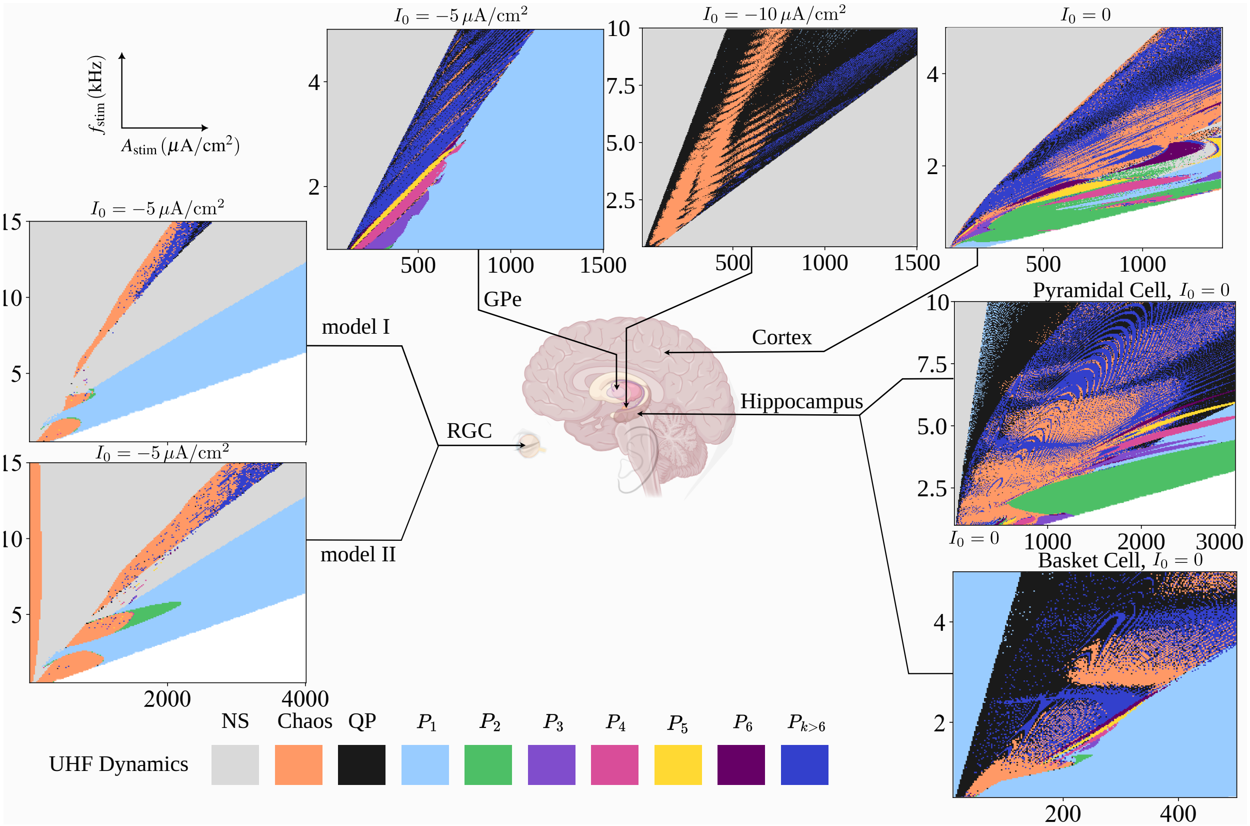


FIG. S15: Functional dynamics map for when mammalian CNS models are stimulated with non-spiking $I_0$ DC values, revealing the presence of NS regions in parameter space.

### S7. Extra average membrane potential displacement maps

Average membrane potential displacement for all other models considered in this work are shown in Fig. S16.

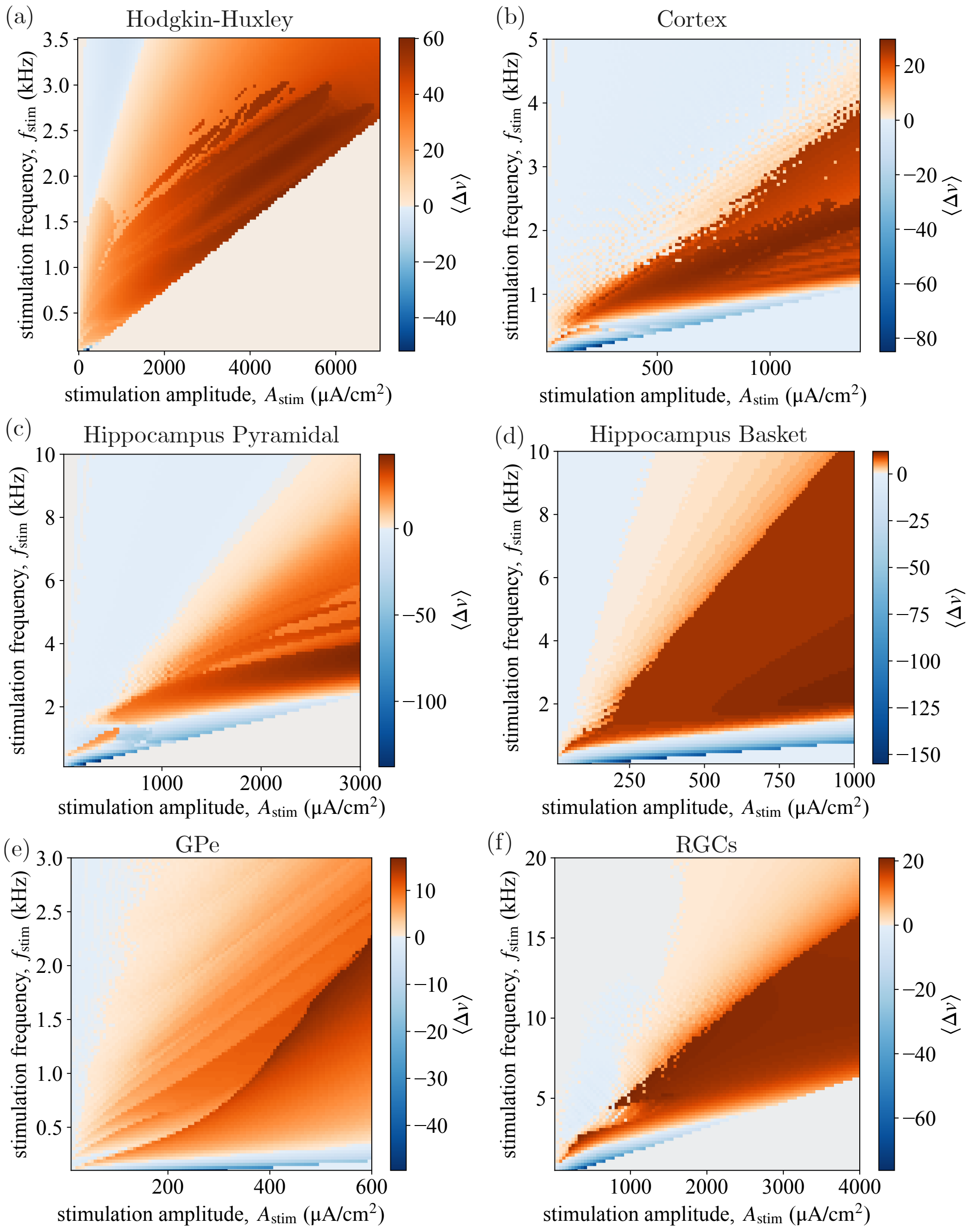


FIG. S16: Extra average membrane potential displacement $\langle \Delta v \rangle$ for (a) Hodgkin Huxley model, (b) Cortex, Hippocampus (c) pyramidal and (d) basket, (e) GPe, and (f) RGCs.

### S8. Additional current equations for mammalian models

For STN and GPe, an additional after-hiperpolarization current $I_{AHP}$, depending on the Ca concentration ($[Ca]$), is called for. It reads

$$I_{AHP} = g_{AHP}\,(v - E_K)\,\frac{[Ca]}{[Ca] + k_1}, \tag{1}$$

where $g_{AHP} = 9\,\mathrm{mS/cm^2}$ ($g_{AHP} = 30\,\mathrm{mS/cm^2}$) and $k_1 = 15$ ($k_1 = 30$) for the STN (GPe) model. Moreover for STN and GPe, $[Ca]$ is modeled as

$$\dot{[Ca]} = -\epsilon\,(I_{Ca} + I_T + k_{Ca}\,[Ca]), \tag{2}$$

with $k_{Ca} = 22.5$ ($k_{Ca} = 20$) and $\epsilon = 3.75 \cdot 10^{-5}\,\mathrm{ms^{-1}}$ ($1 \cdot 10^{-4}\,\mathrm{ms^{-1}}$) for STN (GPe), while $I_T$ and $I_{Ca}$ are currents in the form of Eq. (A11), as defined in Table I, in the main text.

For RGCs, models I and II, the Ca reversal potential $E_{Ca}$ is not a constant, obeying

$$E_{Ca} = \frac{R\,T}{2\,F}\,\ln\left[\frac{[Ca]_e}{[Ca]_i}\right]. \tag{3}$$

The subscripts $e$ and $i$ denote the extra- and intra-cellular spaces, with $R = 8.31451\,\mathrm{J \cdot M^{-1} \cdot K^{-1}}$ the gas constant, $F = 9.6489 \cdot 10^4\,\mathrm{C}$ the Faraday constant and $T$ the temperature in kelvin ($K$). We are taking $T = 308.15\,\mathrm{K}$ ($35^oC$). For both models I and II, the extracellular calcium concentration is fixed $[Ca]_e = 0.0018M$, with the intracellular calcium concentration governed by

$$\dot{[Ca]}_i = -\frac{3\,I_{Ca}}{2\,F\,r}\,\Theta\left[-\frac{3\,I_{Ca}}{2\,F\,r}\right] - \frac{([Ca]_i - Ca_r)}{\tau_{Ca}}, \tag{4}$$

where $r = 0.1\,\mu\mathrm{m}$ is related to the cell surface to volume ratio, $Ca_r = 10^{-7}\,\mathrm{M}$ is the residual calcium concentration inside the cell, $\tau_{Ca} = 1.5\,\mathrm{ms}$ is a proper time constant, $\Theta[\,\cdot\,]$ is the Heaviside step function, and the calcium current $I_{Ca}$ is defined similarly to Equation (A11), as shown in Table I, in the main text. Finally for RGCs, the calcium-activated potassium specific conductance is defined as

$$g_{K,Ca} = \overline{g}_{K,Ca}\,\frac{([Ca]_i/Ca_{\mathrm{diss}})^2}{(1 + ([Ca]_i/Ca_{\mathrm{diss}})^2)}, \tag{5}$$

for $\overline{g}_{K,Ca} = 0.025\,\mathrm{mS/cm^2}$ and $Ca_{\mathrm{diss}} = 10^{-6}\,\mathrm{M}$.

### S9. Derivations of Numerical Lyapunov Exponent Methods

The Lyapunov exponent characterizes the exponential rate of expansion or contraction of state space volumes around a trajectory. Suppose a $N$-dimensional dynamical system

$\dot{x} = F(x, p, t)$, with $x$ the state vector and $p$ a set of parameters. Then, for $N$ linearly independent (LI) small perturbation vectors $\{\delta_1, \ldots, \delta_N\}$ around an initial condition $x_0$, such that $x_0 \to x(t)$ and $x_0 + \delta_0 \to x(t) + \delta(t)$, one shall obtain

$$\delta_i(t) \propto \exp[\lambda_i\, t]\, \delta_i(0), \tag{6}$$

for $t$ small enough and $i = 1, \ldots, N$, Lyapunov exponents $\lambda_1, \ldots, \lambda_N$, ordered as $\lambda_1 \geq \lambda_2 \geq \ldots \geq \lambda_N$. Thus, $\lambda_1$ is the trajectory MLE. Two algorithms are commonly used to obtain the MLE, the computationally costly but reliable QR-method and a faster and direct protocol.

The QR-method computes the full spectrum of exponents, so we just select the largest one. For an initial condition $x_0$ and parameter values $p$, the set of perturbation vectors $\{\delta_1, \ldots, \delta_N\}$ is evolved using the Jacobian $J = dF/dx$ of the dynamical system. The periodic QR-decomposition ensures that perturbations remain orthonormal, preventing the alignment with the fastest expanding direction [1]. While straightforward, this approach demands significant computational time.

A faster method relies only on a single perturbation vector $\delta_0$ around an initial $x_0$. Then, $\delta_0 = [\delta_0/\sqrt{N}, \ldots, \delta_0/\sqrt{N}]^T$ is a N-dimensional vector of isotropic perturbations, with norm $|\delta_0| = \delta_0$. Two trajectories are evolved, departing from $x_0$ and $x_0 + \delta_0$, during a time $t = t_0$, when the Euclidean distance $d$ between both trajectories $T_{x_0}(t_0)$ and $T_{x_0+\delta_0}(t_0)$ escapes the limit $\delta_{\text{down}} < d < \delta_{\text{up}}$. The quantity $\lambda^0 = \ln[d/\delta_0]$ is recorded. The process is repeated for a new initial condition $x_1 = T_{x_0}(t_0)$ and a neighboring $x_1 + \delta_0$, generating $\lambda^1$. The process repeats $M$ times until the total time of integration $T$ is reached. The (fast) MLE is obtained from $\lambda^{\text{fast}} = T^{-1} \sum_i \lambda^i$. This is indeed computationally more efficient than the QR-method, but generally $\lambda^{\text{fast}} \leq \lambda_1$ with $\lambda_1$ from the QR-method. This problem is enhanced in high-dimensional systems.

---


[1] G. Datseris and U. Parlitz, *Nonlinear dynamics: a concise introduction interlaced with code* (Springer Nature, 2022).
